# The regeneration factors ERF114 and ERF115 act as transducers of mechanical cues to developmental pathways

**DOI:** 10.1101/2021.11.29.470368

**Authors:** Balkan Canher, Fien Lanssens, Ai Zhang, Anchal Bisht, Shamik Mazumdar, Jefri Heyman, Frauke Augstein, Sebastian Wolf, Annelie Carlsbecker, Charles W. Melnyk, Lieven De Veylder

## Abstract

Plants show an unparalleled regenerative capacity, allowing them to survive severe stress conditions, such as injury, herbivory attack and harsh weather conditions. This potential not only replenishes tissues and restores damaged organs, but can also give rise to whole plant bodies, highlighting the intertwined nature of development and regeneration. It suggests that regeneration and developmental processes respond to the same upstream signals, but how a cell knows which of the two processes to engage is currently unknown. Here, we demonstrate that next to being regulators of regeneration, ETHYENE RESPONSE FACTOR 114 (ERF114) and ERF115 govern developmental growth in the absence of wounding or injury. Increased ERF114 and ERF115 activity is correlated with enhanced xylem maturation and lateral root formation, whereas their knockout results in a decrease in lateral roots and xylem connectivity following grafting. Moreover, we provide evidence that mechanical cues contribute to *ERF114* and *ERF115* expression in correlation with BZR1 mediated brassinosteroid signaling under both regenerative and developmental conditions. Antagonistically, negative regulation of cell wall extensibility via cell wall-associated mechanosensory FERONIA signaling suppresses their expression under both conditions. Our data suggest a molecular framework in which mechanical perturbations too great to be compensated by adaptive cell wall remodeling results in strong *ERF114* and *ERF115* expression, switching their role from developmental to regenerative regulators.

**One-Sentence Summary:** Mechanical cues drive *ERF114/ERF115* expression during lateral root development as well as regeneration.

## INTRODUCTION

While inherited genetic and molecular information in plants and animals orchestrate the unfolding of developmental processes with high precision, strategies to cope with unexpected loss of cells or organs through wounding are essential for survival and continuity of any species. Due to their immobile nature, plants have developed mechanisms that allow the organism to instantly detect the nature and extent of injury and activate the precise regenerative mechanisms needed for each of the numerous scenarios of injury. Stem cell death, meristem incision or targeted wounding such as loss of a few cells upon laser ablation, locally reactivate the developmental signaling pathways that allow replacement of the lost cells in a precise and functional manner (Birnbaum and Sanchez Alvarado, 2008; Canher et al., 2020; Melnyk et al., 2015; Savatin et al., 2014). Interestingly, regenerative signaling networks governing callus formation are activated during lateral root formation, suggesting that regeneration and development has been described as two sides of the same coin (Perianez-Rodriguez et al., 2014). An outstanding research question is how can the organism differentiate whether activation of a developmental signaling pathway is needed as a regular part of development or in a regenerative context?

The ERF subfamily X transcription factors (ERF108-115) has been suggested to play an emerging role in diverse developmental and regenerative processes (Heyman et al., 2018). ERF115 was initially identified as a driver of quiescent center stem cell divisions during development (Heyman et al., 2013). Since the discovery that death of a single cell triggers *ERF115* expression in neighboring cells and subsequently pushes them to initiate a cell division program, it has been studied extensively within the context of wounding and regeneration (Heyman et al., 2016; Hoermayer et al., 2020; Marhava et al., 2019; Zhou et al., 2019). Expression of *ERF115* after DNA damage-induced stem cell death was found the act synergistically with wound induced accumulation of auxin, which in turn allows the damaged cells to be replaced by new cells (Canher *et al*., 2020). The synergistic effect of ERF115 on auxin signaling was reported to be due to activation of *MONOPTEROS* (*MP*), a member of the *AUXIN RESPONSE FACTOR* (*ARF)* family. *MP* has been shown to be essential for many developmental processes including maintenance and formation of meristems, embryonic root formation, flower formation and vascular development (Bhatia et al., 2016; Jürgens, 1993; Luo et al., 2018; Przemeck et al., 1996). Similar to its role in primary meristems, *MP* mediated auxin signaling and auxin transport tightly controls virtually every stage of LR development (De Smet et al., 2010).

ERF114, the closest homolog of ERF115, is also known as ERF BUD ENHANCER (EBE) because of the observed increased axillary bud outgrowth upon its overexpression (Mehrnia et al., 2013). Not only is its expression strongly induced following wounding and coincides with callus formation at the cut sites, a prominent feature observed upon *ERF114* overexpression is neoplasia in the form of tissue that is similar to green callus, and it is often produced at wound sites. Moreover, it has been observed that *ERF114* overexpression tissue explants display increased rates of callus production when cultured on callus-inducing medium (Mehrnia *et al*., 2013) Even though the involvement of ERF subfamily X transcription factors in wounding and regeneration is well established, the mechanism leading to their activation upon wounding has not been agreed upon. Activation of a jasmonate signaling network acting in synergy with auxin was reported to cause *ERF115* induction upon wounding or nematode infection

(Zhou *et al*., 2019). Furthermore, reactive oxygen species were shown to control the expression of *ERF114* and *ERF115* for maintaining the stem cell division and differentiation balance (Kong et al., 2018). Hoermayer *et al*. demonstrated that following targeted cell death by means of laser ablation, turgor driven expansion of the adjacent cell was a prerequisite to *ERF115* activation. Furthermore, damaging of the cell wall without killing the cell was sufficient for *ERF115* induction in the presence of auxin (Hoermayer *et al*., 2020). These data indicated a potential role for cell wall integrity signaling and acute mechanical stress in wound-induced *ERF115* activation.

One of the main drivers of plant growth and cell expansion is the dynamic interplay between intrinsic and extrinsic mechanical forces where the direction of growth is determined by the extensibility of the cell wall. Plant cell walls are mainly made up of cellulose fibers embedded in a pectin-cellulose-hemicellulose carbohydrate matrix (Hofte and Voxeur, 2017). The extensibility of the cell wall is determined by the type and extent of crosslinks between the aforementioned components. During development the internal and external forces experienced by the cell wall are transmitted into downstream cellular signaling pathways through a variety of cell membrane localized receptors, directly linked with cell wall components. This provides the input for the cell to expand or divide, accompanied by remodeling of the cell wall to maintain homeostasis (Hofte, 2015). One of the best studied family of cell wall surveillance proteins is plant malectin-like receptor kinases, also known as CATHARANTHUS ROSEUS RECEPTOR-LIKE KINASE 1-LIKE PROTEINS (CrRLK1Ls) (Franck et al., 2018). FERONIA (FER), a well-studied member of CrRLK1Ls family, has been shown to bind pectin *in vitro*, suggesting its association with the cell wall (Feng et al., 2018). Mutants lacking FER activity display accelerated growth, increased strain rate experienced by the root and hypersensitivity to abiotic stresses (Feng *et al*., 2018; Shih et al., 2014). Based on these observations, FER is proposed to sense changes in the mechanical equilibrium of cell walls and negatively regulate cell wall extensibility to maintain cell wall homeostasis (Hofte, 2015). This inhibitory regulation of cell expansion contrasts with the positive regulation of cell expansion and elongation mediated by the brassinosteroid (BR) hormone signaling pathway. BR synthesis occurs locally in rapidly elongating cells and acts in a paracrine fashion through its receptor BRASSINOSTEROID INSENSITIVE 1 (BRI1) (Vukasinovic et al., 2021). The subsequent activation of transcription factor BRASSINAZOLE RESISTANT 1 (BZR1) has been shown to activate a plethora of adaptive downstream signaling cascades enriched in genes responsible for cell wall modification (Sun et al., 2010). Perturbation of cell wall integrity through pectin methyl esterase inhibition has been shown to activate BR signaling leading to adaptive changes in growth and cell wall remodeling (Wolf et al., 2012b; Wolf et al., 2014).

In this study, we demonstrate that ERF115 and its closest homolog ERF114 regulate xylem tissue maturation and lateral root (LR) development during regular growth. Moreover, using the *feronia* cell wall integrity receptor mutant, we demonstrate that mechanical strains contribute to *ERF114* and *ERF115* expression in both developmental and regeneration contexts by activation of BR signaling. The data highlight an intimate relationship between development and regeneration that is under control of ERF114 and ERF115.

## RESULTS

### ERF114 acts redundantly with ERF115 during regeneration following root tip removal

*ERF114*, the closest homolog of *ERF115*, is also known as *ERF BUD ENHANCER* (*EBE*) because of the observed increased axillary bud outgrowth upon its overexpression (Mehrnia *et al*., 2013). Its expression is strongly induced following wounding and coincides with callus formation at the cut sites, similar to *ERF115* (Heyman *et al*., 2016). However, the importance of ERF114 for the reconstitution of an organized root meristem following wounding has not been explored. We generated a reporter line in which the NLS-GFP/GUS fusion protein is driven by the *ERF114* promoter to study its root-specific expression under control and wounding conditions. Confocal imaging revelated that *ERF114* was induced strongly 5 h after removal of the distal root tip (hours post cut; hpc) and its expression diminished as a new root tip was formed at 72 hpc (Fig. 1A). Similarly, stem cell death caused by 24-h treatment with the radiomimetic drug bleomycin (BLM) evoked a strong *ERF114* induction around dead stem cells (Fig. 1B). Seeing the similarity in transcriptional activation of *ERF114* and *ERF115* following tissue-damaging conditions, we suspected a functional redundancy between these two close homologs (Canher *et al*., 2020; Heyman *et al*., 2016). Therefore, we created *erf114* single and *erf114 erf115* double mutants via CRISPR/Cas9-mediated mutagenesis to overcome genetic redundancy. Next, we evaluated their ability to regenerate a de novo root tip 72 h after stringent root tip cutting, being the removal of the 200µm distal root tip. As previously observed (Heyman *et al*., 2016), the dominant-negative *ERF115^SRDX^* overexpression line and *erf115* single mutants displayed a lower regeneration frequency compared to the wild type, being 1%, 31% and 52% respectively (Fig. 1C). *erf114* single mutants displayed a slightly lower regeneration frequency of 44%, which was not statistically significant compared to the wild type. However, additional mutation of *erf114* in the *erf115* mutant background significantly reduced the regeneration frequency compared to the wild type, being between that of the *ERF115^SRDX^* and *erf115* single mutants (15%) (Fig. 1C). Taken together, these results suggested that ERF114 acts redundantly with ERF115 during root meristem regeneration.

**Fig. 1.**
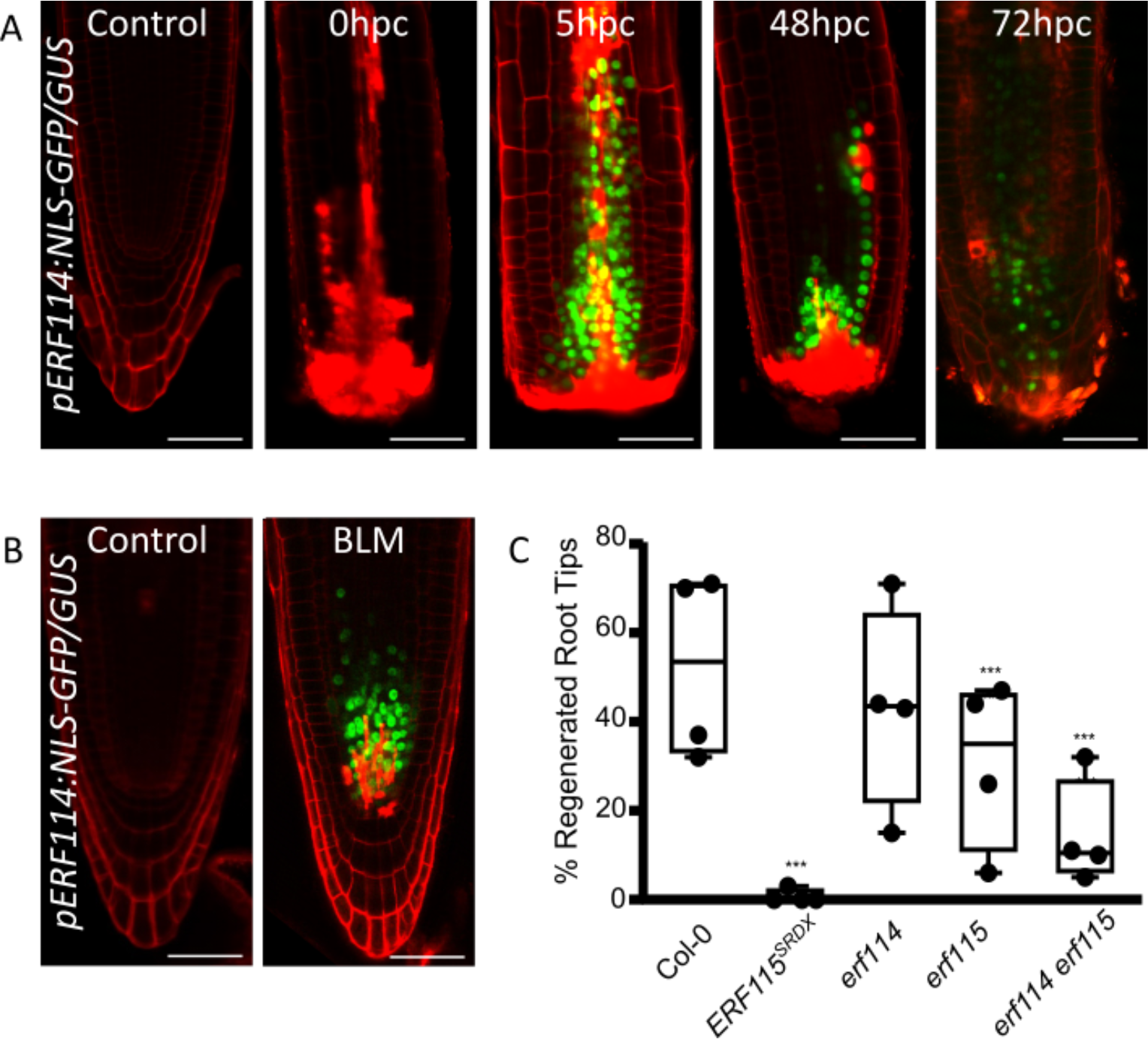
ERF114 and ERF115 redundantly control root tip regeneration. **A-B**, Confocal images of *pERF114:GFP-NLS/GUS* root tips (red) during regeneration from root tip excision (**A**) or after 24h bleomycin (BLM) treatment (**B**). (hpc=hours post cutting). Cell walls and dead cells are counterstained (red) with propidium Iodide (PI) Scalebars = 50 μm **C,** Percentage of regenerated root tips after stringent root tip excision. Error bars indicate standard error. Significance was calculated based on Fisher’s exact test. (*** p<0.001)

### *ERF114* overexpression enhances sensitivity to auxin accumulation

ERF115 has previously been shown to act upstream of MONOPTEROS (MP), a major regulator of various auxin-mediated development processes (Canher *et al*., 2020). In addition, overexpression of *ERF115* was shown to confer increased sensitivity to auxin accumulation in the meristem caused by chemical inhibition of polar auxin transport by naphtylphthalamic acid (NPA). Following two weeks of growth on NPA-containing medium, *ERF115^OE^* seedlings display enhanced vascular diameter as well as an expanded columella domain, as indicated by the presence of starch granules (Canher *et al*., 2020). We aimed to investigate if *ERF114* overexpression would result in a similar sensitivity to auxin accumulation. After 5 days of growth on NPA, both *ERF114^OE^* and *ERF115^OE^* seedlings displayed enhanced expansion of the columella domain, compared to the wild type (Fig. 2A). Likewise, after 14 days of growth on NPA, enhanced vascular diameter was observed both in *ERF114^OE^* and *ERF115^OE^* seedlings compared to the wild type (Fig. 2B,C). Besides the previously reported effects on columella domain and vascular expansion, we observed that both *ERF114^OE^* and *ERF115^OE^* seedlings grown on NPA displayed ectopic xylem formation likely because of transdifferentiation of cambial cells. In contrast to the absence of differentiated xylem cell files in wild-type roots, both *ERF114^OE^* and *ERF115^OE^* seedlings displayed multiple xylem strands at 200-250µM from the root tips (Fig. 2B,C). These data suggest that *ERF114* and *ERF115* might contribute to xylem formation in response to increased auxin signaling.

**Fig. 2.**
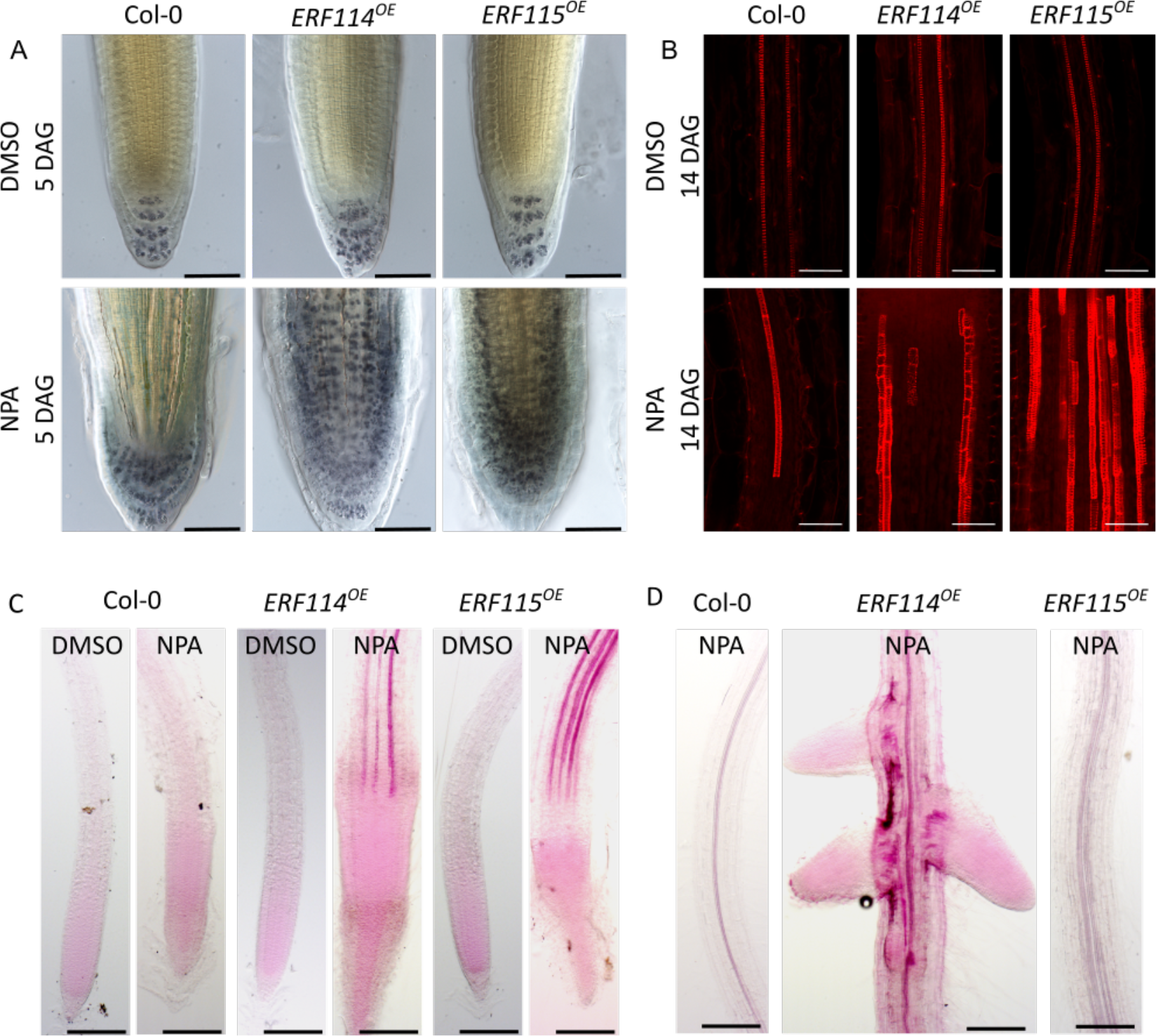
ERF114 and ERF115 enhance auxin-driven xylem maturation and lateral root initiation. A, **A**, Light microscopy images of lugol-stained root tips grown on DMSO or 10µM NPA for 5 days. DAG, days after germination. **B-D,** Confocal (**B**) and light (**C,D**) microscopy images of basic fuchsin stained roots grown on DMSO or 10µM NPA for 14 days. Highly stained (red) structures indicate multiple strands of lignified xylem cells. (**D**) Lack of lateral roots in NPA grown Col-0 and *ERF115^OE^* seedlings in contrast to abnormal lateral root formations observed in *ERF114^OE^* seedlings grown on NPA Scale bars = 50 μm.

Transdifferentiation of cambial cells into xylem vessels is an essential part of the regenerative process during grafting for re-establishment of water and nutrient transport between the rootstock and scion. Furthermore, hypocotyl cutting has been associated with an accumulation of auxin (Melnyk et al., 2018). Due to the well-established role of ERF115 in various regenerative processes, we hypothesized that ERF114 and ERF115 might also be involved in xylem connectivity upon hypocotyl grafting. Confocal imaging revealed no clear morphological differences at xylem reconnection sites between the analyzed genotypes as indicated by basic fuchsin staining (Fig. 3A). Therefore, we performed grafting assays and quantified the percentage of grafts that had successful xylem reconnection based on the presence of fluorescence in the scion following 5,6- carboxyfluoescein diacetate (CFDA) addition of the rootstock. The xylem reconnection rate did not change substantially for seedlings overexpressing *ERF114* (94%) while there was a significant but small increase in those overexpressing *ERF115* (97%, Fig. 3B), probably due to the already high reconnection rate for wild-type plants (90%). In contrast, the *erf114 erf115* double mutant and *ERF115^SRDX^* dominant-negative lines had a substantially decreased xylem reconnection rate (53%, p<0.01 and 19%, p<0.01, respectively). These results demonstrate that *ERF114* and *ERF115* play an important role in establishing xylem reconnection between rootstock and scion following grafting. Considering that high levels of *ERF114* and *ERF115* expression is generally observed in wounded tissue, the observed decreased rate of xylem reconnection during hypocotyl grafting in *erf114 erf115* mutant supports the link with ERF-mediated regeneration and xylem differentiation.

**Fig. 3.**
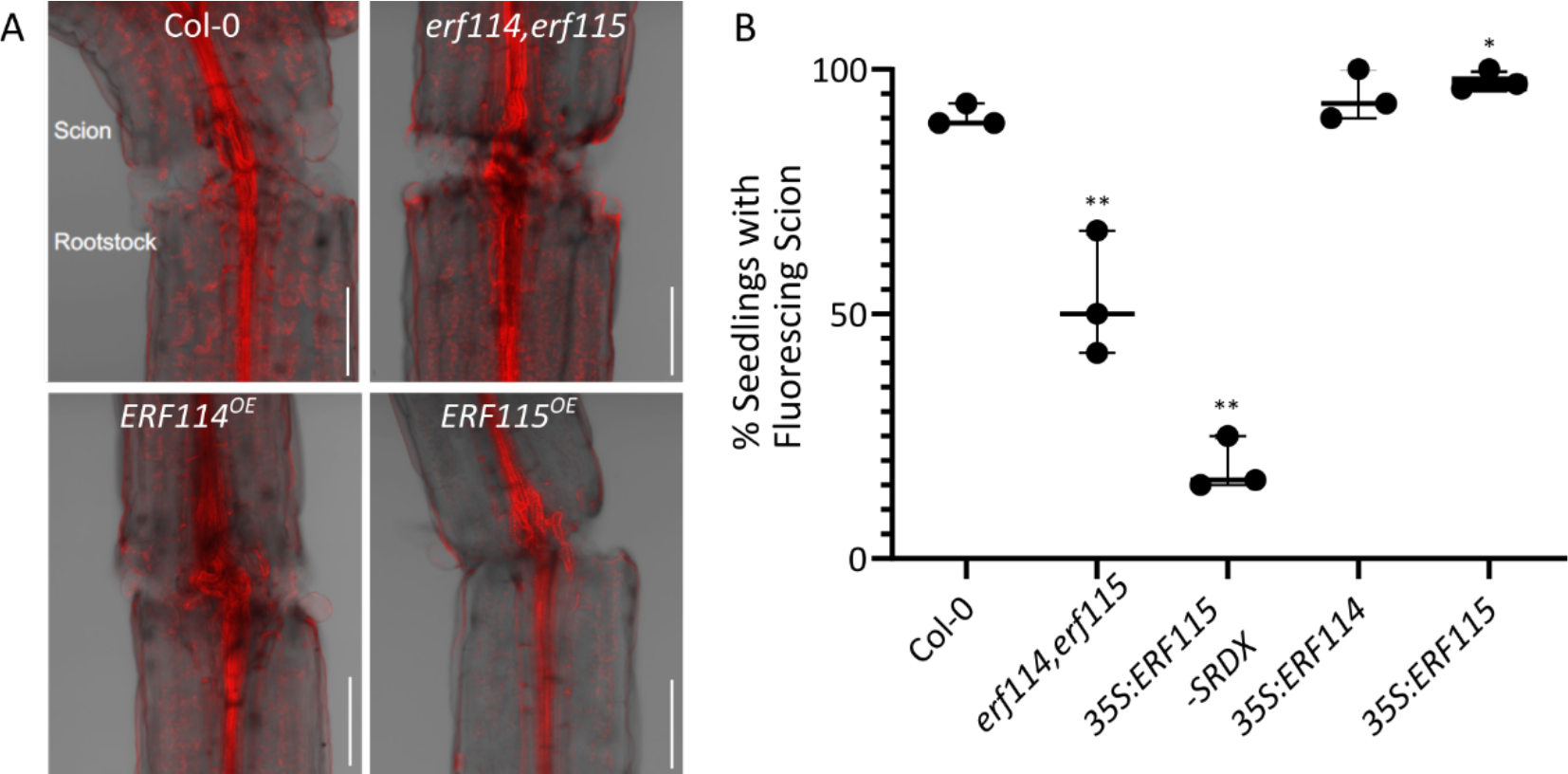
ERF114 and ERF115 are required for xylem connectivity following grafting. (A) Confocal images of pseudo schiff Propidium Iodide (mPS-PI) stained hypocotyls marking the xylem strands. Scalebar = 50 μm. (B) Xylem reconnection rates 7 days after grafting as indicated by fluorescing scion. Statistical analysis of differences compared to Col-0 were performed using Logistic Regression modeling. Odds ratios and p-values were calculated using Fisher’s exact test (*p<0.05, **p<0,01, ***p<0.001). Scale bars = 50 μm.

While these data suggest a similar functionality between *ERF114* and *ERF115* during regeneration, long-term NPA treatment revealed potential differences in functionality. Growth in the presence of NPA has been found to inhibit LR primordium (LRP) formation and outgrowth, due to the essential role of polar auxin transport in the formation of lateral organs (Casimiro et al., 2001). In accordance with these reports, 14-day-old wild-type seedlings grown on 10 µM NPA produced no LRs (Fig. 2D). Similarly, no LRPs were observed in *ERF115^OE^* seedlings. Surprisingly, *ERF114^OE^* seedlings grown on NPA displayed developed LRs (Fig. 2D), demonstrating that high *ERF114* levels allow to initiate lateral roots regardless of auxin transport inhibition.

### Both ERF114 and ERF115 are part of the developmental LR formation program

ERF115 has been shown to act as an activator of MP, a master regulator for a plethora of developmental auxin mediated developmental processes including LR formation and vascular development (Bhatia *et al*., 2016; Canher *et al*., 2020; De Smet, 2010; Przemeck *et al*., 1996). Due to the formation of LRs on NPA-treated *ERF114^OE^* seedlings, we investigated if *ERF114* and *ERF115* might be involved in LR formation under control conditions. F1 seedlings resulting from the cross between *MP:MP-GFP* and *ERF115^OE^* had higher MP levels in various stages of LR formation (Fig. 4A), suggesting that ERF115 activity might promote LR development under regular growth conditions.

**Fig. 4.**
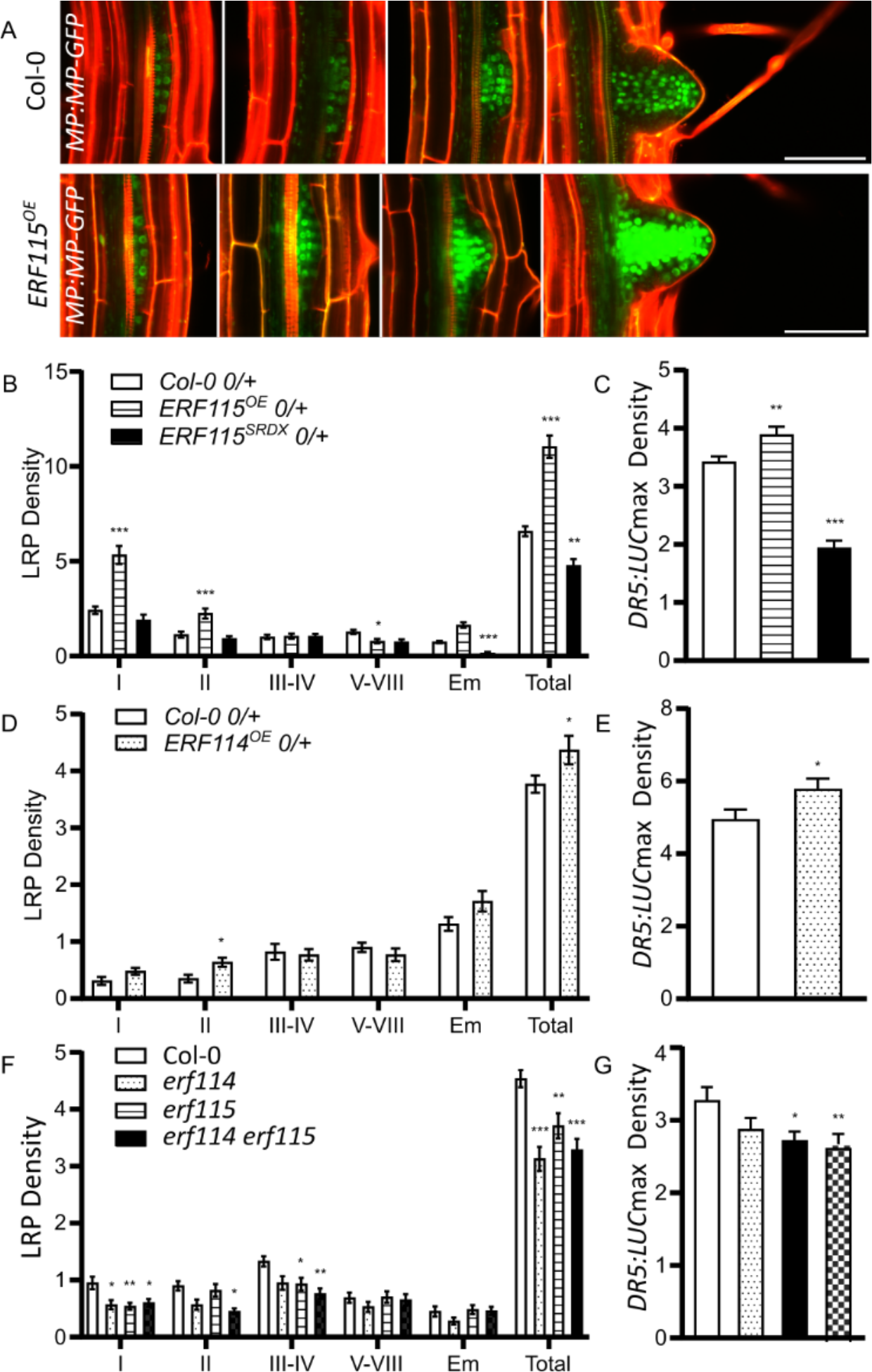
ERF114 and ERF115 control lateral root primordia initiation. **A,** Confocal images off PI-stained lateral root primordia (LRP) showing MP:MP-GFP in wild type (Col-0) and *ERF115^OE^*. backgrounds. Scale Bars = 50 μM **B-E,** LR (**B,D**) and *DR5:LUC* maxima densities (**C,E**) of indicated genotypes crossed with *DR5:LUC* seedlings in the F1 generation (indicated as 0/+). **F-G,** LRP (**F**) and DR5:LUC maxima densities (B) of homozygous *erf114*, *erf115* and *erf114 erf115* seedlings carrying *DR5:LUC* construct (D-E) (*p<0.05, **p<0,01, ***p<0.001 mixed model procedure was used to calculate significance values for pairwise comparisons with the wild type by least squares post-hoc tests and corrected for multiple testing using the Bonferroni Method )

*DR5:LUCIFERASE* (*DR5:LUC*) reporter is commonly used to visualize the LRP sites marked by auxin maxima (Moreno-Risueno et al., 2010).To examine if *ERF114* and *ERF115* control LRP initiation, we crossed the *DR5:LUC* reporter with the *ERF115^OE^* and *ERF115^SRDX^* overexpression lines and analyzed the auxin maxima density (*DR5:LUCmax*) in the F1 generation from luciferase luminescence images (Supplementary Fig. 1A,B). LRP densities were also determined from differential interference contrast images after clearing as previously described (Malamy, 1997). The *ERF115^OE^* hemizygous seedlings showed significantly increased LRP (Fig. 4B) and *DR5:LUCmax* (Fig. 4C) densities compared to the wild type, in contrast to the *ERF115^SRDX^* seedlings showing a decrease in both. Similarly to *ERF115^OE^*, *ERF114^OE^* seedlings had increased LR and *DR5:LUC* maxima densities (Fig. 4D,E).

Next, we investigated how *ERF114* and *ERF115* loss of function would impact LR development. To this end, we introduced the *DR5:LUC* reporter into *erf114* and *erf115* single and *erf114 erf115* double backgrounds. We noticed that the *erf114* and *erf115* single mutants and the *erf114 erf115* double mutants displayed a significantly reduced LR density compared to wild-type plants (Fig. 4F). Luminescence imaging of *DR5:LUC* showed that *erf115* single and *erf114 erf115* double mutants displayed a significant reduction in *DR5:LUC* maxima density compared to the wild type (Fig. 4G). The *erf114* single mutant also showed a reduced *DR5:LUC* maxima density but the difference was not statistically significant.

To confirm the involvement of *ERF115* in the production of auxin maxima, we rotated the plates on which the *DR5:LUC ERF115^SRDX^* seedlings were growing for 90°, a process known to induce a gravitropic curvature in the roots accompanied by the accumulation of auxin at the bend site (Lucas et al., 2008). Luminescence time-lapse imaging after 90° rotation indicated that a significantly reduced number of auxin maxima formed at the bend site of *DR5:LUC ERF115^SRDX^* roots compared to the wild type (Supplementary Fig. 1C), further suggesting the involvement of ERF115 in the formation of bending-induced auxin maxima. Taken together, these data suggest that next to wound induced regeneration, both ERF114 and ERF115 are involved in LR formation and development.

### Spatial-temporal control of *ERF114* and *ERF115* expression reveals correlation between protoxylem maturation and LR initiation

Despite the data indicating a similarity between ERF114 and ERF115 activity and LR development, expression of these transcription factor genes in LR primordia has thus far not been reported, which prompted us to investigate potential LR-specific *ERF114* and *ERF115* expression in detail. Confocal imaging of *pERF115:NLS-GFP/GUS* seedlings, grown under control conditions, showed a relatively weak (requiring high laser power) and sporadic expression of *ERF115* in protoxylem cells (Fig. 5A), in line with a previous study (Zhou *et al*., 2019). *PASPA3* expression has been shown to mark maturing protoxylem cells undergoing programmed cell death (Fendrych et al., 2014). Time-lapse imaging of a *pPASPA3:tdTOMATO-NLS pERF115:NLS-GFP/GUS* dual reporter line showed that *ERF115* expression overlapped with that of *PASPA*3, suggesting that *ERF115* expression correlates with the maturation of protoxylem cells (Fig. 5D, Supplementary Movie 1). The sudden disappearance of both fluorescent signals in the older protoxylem cells (as indicated by the disappearance of arrows in Fig. 5D and clearly shown in Supplementary Movie 1) likely results from nuclear disintegration in the last stages of programmed cell death. However, no *ERF115* expression could be detected in LRP. To investigate if *ERF115* expression was also lacking in uninitiated LRs named prebranch sites, we introduced the *DR5:RFP* reporter, which is activated prior to LR initiation (Moller et al., 2017), into the *ERF115* reporter line. Time-lapse imaging using the *pERF115:GFP-NLS pDR5:RFP* dual reporter line confirmed that the prebranch sites marked by the *DR5:RFP* signal did not harbor a detectable *ERF115:GFP-NLS* signal (Supplementary Movie 2). However, we noticed that the *ERF115:GFP-NLS* signal in the protoxylem had a tendency to disappear shortly after the *DR5:RFP* signal started accumulating in the LR founder cells, which subsequently forms the LRP (Supplementary Movie 2). To investigate this correlation in more detail, time-lapse imaging was performed after rotating the seedlings 90°C to synchronously induce LR formation. The *DR5:RFP* signal appeared in the founder cell 1.6 ± 0.7 h (n=6) before the disappearance of the *ERF115:GFP-NLS* signal in the adjacent protoxylem, suggesting a temporal correlation. Likewise, *pDR5:VENUS-NLS pPASPA3:tdTOMATO-NLS* time-lapse imaging revealed that the *DR5:VENUS-NLS* signal in the founder cell appeared on average 1.1 ± 0.5 h (n=7) before the *pPASPA3:tdTomato-NLS* signal disappeared in the adjacent protoxylem (Supplementary Movie 3, Supplementary Fig. 2). These results suggest that while *ERF115* is not expressed in the LR founder cell, there is a temporal correlation between the activation of auxin signaling in the founder cell and the nuclear degradation of adjacent protoxylem cells expressing *ERF115*.

**Fig. 5.**
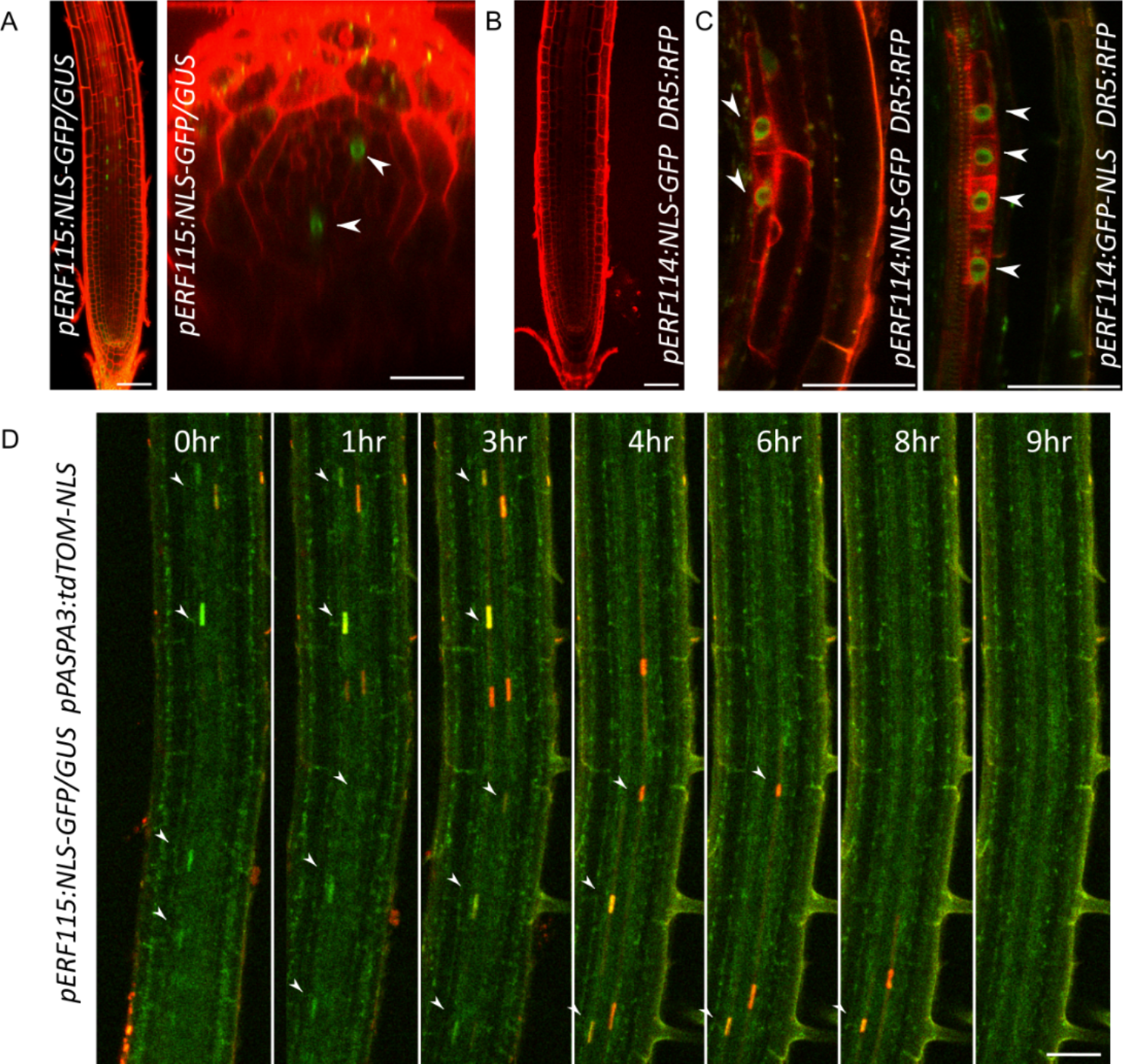
Developmental controlled expression of *ERF114* and *ERF115*. **A**, Confocal image of *pERF115:GFP-NLS/GUS* root and cross section image showing protoxylem specific *ERF115* expression (indicated by arrowheads). **B,** Confocal image of *pERF114:GFP-NLS/GUS* root. **C**, Confocal images of *pERF114:GFP-NLS/GUS DR5:RFP* stained with PI showing *ERF114* expression in the LR founder cells (indicated by arrowheads). **D,** Stills from time lapse imaging of the *pERF115:GFP-NLS/GUS pPASPA3:tdTomato-NLS* dual reporter during protoxylem maturation (see also Supplemental Movie 1). White arrowheads point to *pERF115:GFP- NLS/GUS* positive protoxylem nuclei. Yellow colored nuclei contain both *pERF115:GFP- NLS/GUS (green)* and *pPASPA3:tdTomato-NLS* (red) signals. Scale bars = 50 μm.

Differently from *ERF115*, expression of *ERF114* could be detected in pericycle cells while being absent in the protoxylem (Fig. 5A,B). Expression of *ERF114* in the pericycle was accompanied by a *DR5:RFP* signal, which marks the LR founder cell prior to initiation (Fig. 5C). *ERF114* expression in the founder cell preceded the first appearance of the *DR5:RFP* signal by 1.4 ± 1.1 h (n=6) (Supplementary Movie 4, Supplementary Fig. 3). Similarly, *ERF114* expression in the founder cell preceded the protoxylem cell death by 2.75 ± 0.5 h (n=6).

### Cell wall damage, mechanical stress and endodermal cell removal induce *ERF114* expression

Hoermayer *et al*. recently demonstrated that following laser-assisted cell ablation, cell expansion is necessary for the induction of *ERF115*. Moreover, damaging the cell wall without causing cell death in the presence of auxin was sufficient for robust induction of *ERF115* (Hoermayer *et al*., 2020). Based on these observations, we hypothesized that the expression of *ERF114* and *ERF115* during the process of LR formation might be influenced by mechanosensitive cues or cell wall stresses. To test this hypothesis, we damaged the cell wall at the intersection of two immature protoxylem cells using low-intensity laser exposure without causing cell death, as indicated by lack of intense PI staining (red) (Fig. 6A). Protoxylem ablation in the *pERF114:GFP-NLS/GUS* background was followed by rapid induction of *ERF114* in the adjacent pericycle cells, suggesting that cell wall damage signals coming from the neighboring protoxylem indeed can induce pericycle-specific *ERF114* expression (Fig. 6A, Supplementary Movie 5). Previously, it has been proposed that endodermal cells act as inhibitors of LR development and that laser-assisted endodermal cell ablation (ECA) triggers periclinal divisions in the underlying pericycle cells. Mutants impaired in auxin signaling and transport still display periclinal divisions upon ECA, albeit in a reduced frequency, hinting towards the involvement of additional processes independent of auxin signaling (Marhavy et al., 2016). To check whether *ERF114* might be involved in this process, we performed ECA using a high laser intensity. ECA triggered strong and rapid *ERF114* induction in the pericycle followed by periclinal and anticlinal divisions (Fig. 6B). While it is tempting to speculate about a potential causative link between protoxylem cell death and *ERF114* and *ERF115* expression, maturation of the protoxylem has not been found to be necessary for LR initiation (Parizot et al., 2008). Therefore, it is unlikely that the cell death itself is triggering *ERF114* and *ERF115* expression in LR founder cells and the protoxylem, respectively. Transient mechanical bending of roots has been shown to be sufficient to induce an LR even in mutants with severely impaired auxin signaling such as *tir-1* and *arf7 ar19* (Ditengou et al., 2008). Furthermore, LRs have been shown to be located at the sites of curvature resulting from auxin-dependent root waving (De Smet et al., 2007). Therefore, we hypothesized that instead of the cell death, the localized mechanical strains associated with natural curvature of the root growth might be developmental triggers for *ERF114* and *ERF115*. To test this hypothesis, we applied mechanical stress by manual bending the root tip and releasing it, as described previously (Ditengou *et al*., 2008). Absence of intense PI staining (cyan) suggested that no cell death occurred during the process (Fig. 6C, Supplementary Movie 6). Manual root bending triggered strong *ERF114* expression in multiple pericycle cells along the bent region, followed by LR primordia initiation in a subset of *ERF114* positive cells carrying a strong *DR5:RFP* signal (Fig. 6C, Supplementary Movie 6). Similarly, manual bending also caused rapid induction of *ERF115* but in tissues underlying the LR primordia carrying the *DR5:RFP* signal (Supplementary Movie 7). These data demonstrate that mechanical stress is sufficient to trigger both *ERF114* and *ERF115* expression in the pericycle and vasculature, respectively.

**Fig. 6.**
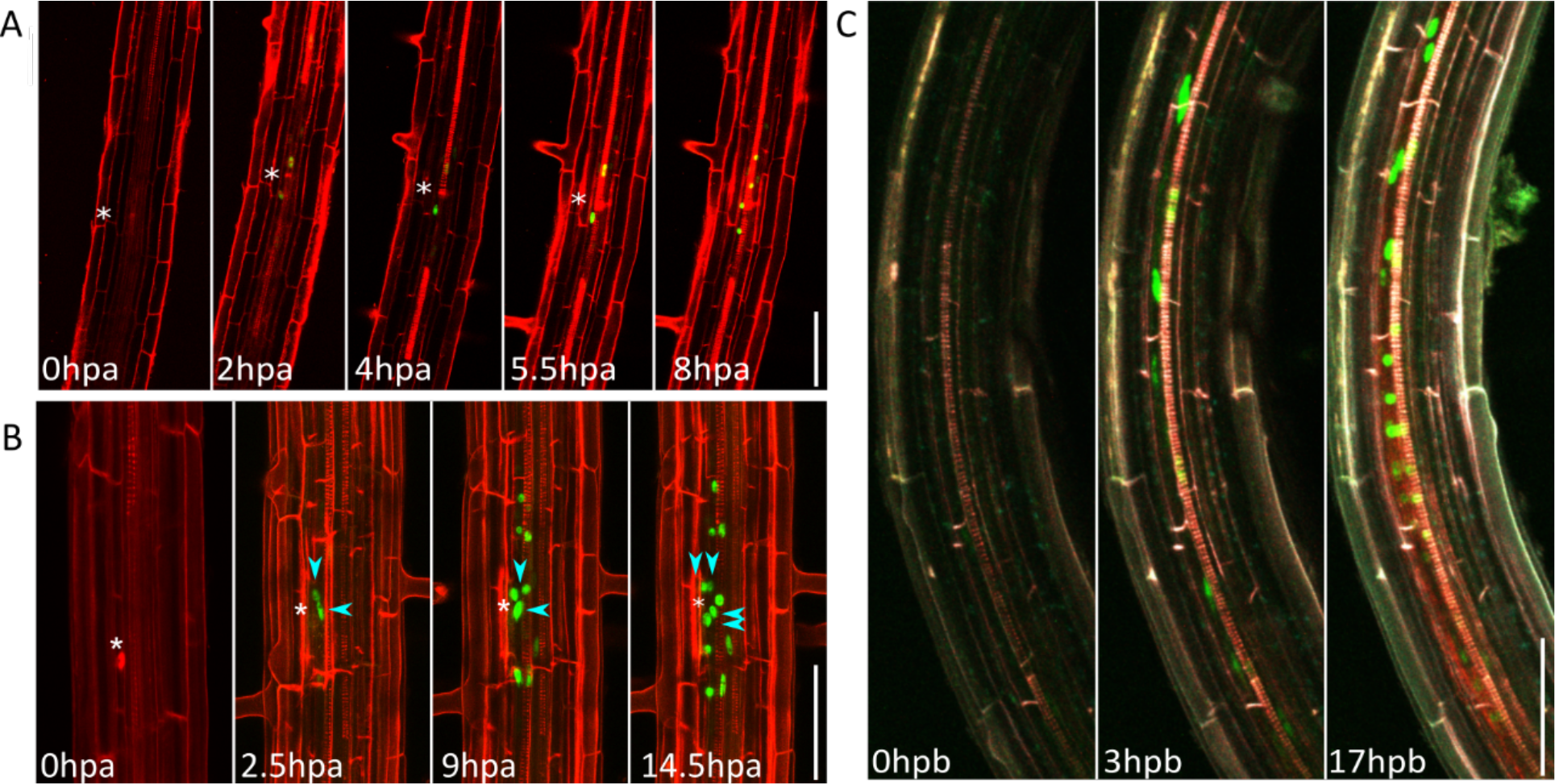
*ERF114* responds to cell wall damage and mechanical stress. **A-B,** Stills from time lapse imaging of PI stained (red) *pERF114:NLS-GFP/GUS* (green) seedlings after laser induced damaging of on immature protoxylem cell wall (Supplementary Movie 5) (**A**) or after endodermal cell ablation. (**B**). Asterisks mark the point of laser exposure in the cell wall boundary between two protoxylem cells and ablated endodermal cell. Blue arrowheads indicate periclinal divided nuclei. **C,** Stills from time lapse imaging (Supplemental Movie 6) of PI-stained (cyan) *pERF114:NLS-GFP/GUS* (green) p*DR5:RFP* (red) seedling after mechanical stress induced by root bending. hpa: hours post ablation hpb: hours post bending. Scale bars = 50 μm

### The cell wall-associated mechanosensor FERONIA receptor kinase limits developmental and wound-induced *ERF114* and *ERF115* expression

Cell wall damage signaling has been reported to activate BR hormone signaling that in turn modifies the cell wall architecture to increase cell wall extensibility (Wolf *et al*., 2012b). Following DNA damage-induced stem cell death caused by BLM, cell expansion of neighboring cells has been shown to be necessary for *ERF115* activation (Hoermayer *et al*., 2020). Since BZR1, one of the main regulators of BR signaling was shown to bind directly to the *ERF115* promoter, and BL treatment enhanced its expression levels, we tested if it is activated upon BLM treatment in expanding cells neighboring the dead ones (Heyman *et al*., 2013; Lee et al., 2015). Confocal imaging of the BR signaling reporter *pBZR1:BZR1-GFP* suggested that BLM-induced stem cell death results in an accumulation of BZR1 around the dead cells (Fig. 7A), in agreement with the previously reported activation of wound-induced BR signaling (Wolf et al., 2012a). We reasoned that activation of BZR1 during the LR formation could be indicative of perceived cell wall stress. Time-lapse imaging using the *pBZR1:BZR1-GFP* reporter line indicated that nuclear BZR1 accumulation can be observed in the LR founder cells and initiated primordia as marked by *DR5:RFP* accumulation (Fig. 7B).

**Fig. 7.**
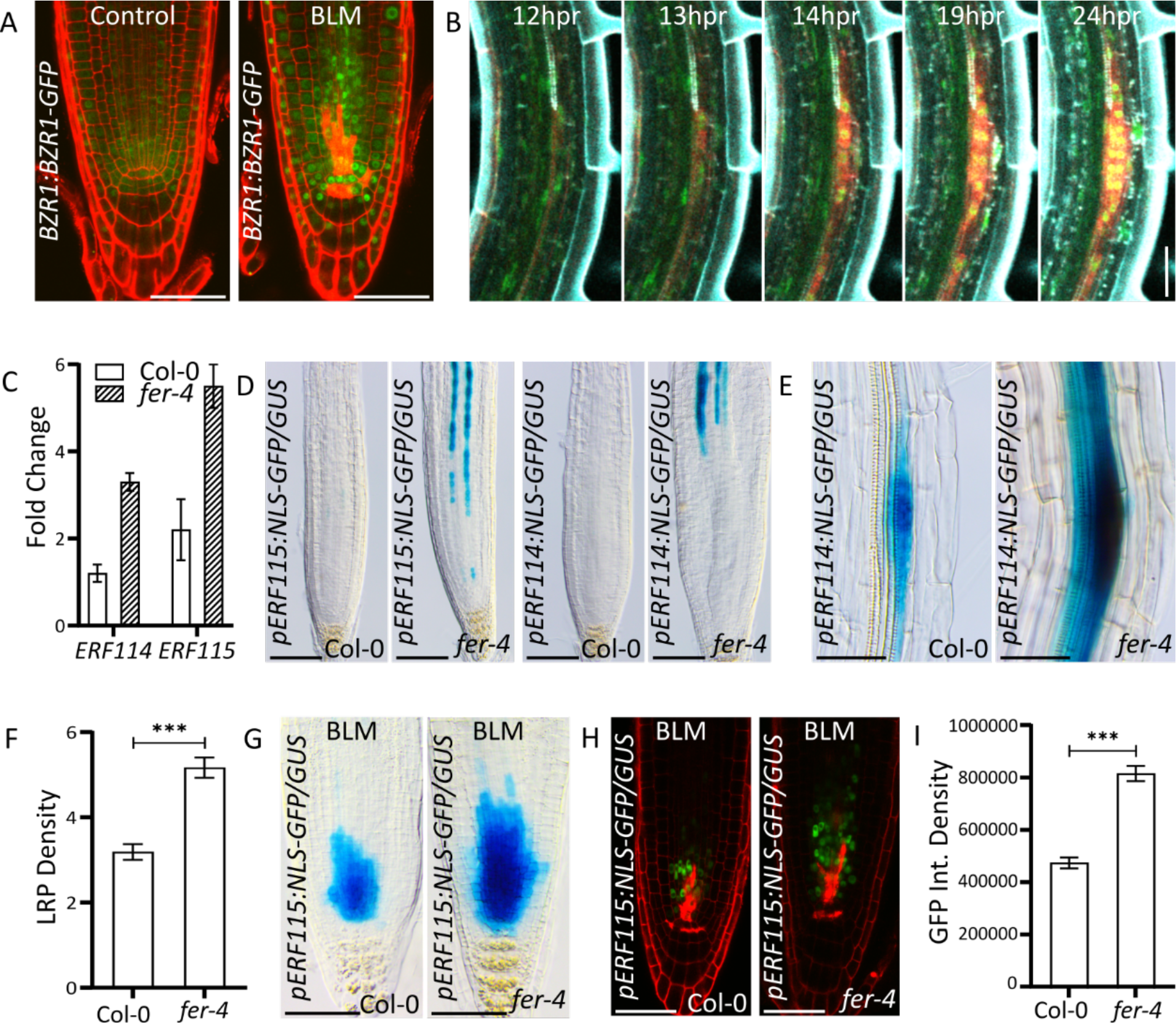
*ERF114* and *ERF115* respond to cell wall mechanosensory signals. **A,** *pBZR1:BZR1-GFP* expression under control conditions and after 24h BLM treatment (Intense PI staining (red) indicates cell death) **B,** Time lapse imaging of PI stained (cyan) BZR1:BZR1-GFP (green) DR5:RFP (red) during LR formation. **C,** Fold changes in *ERF114* and *ERF115* expression in wild type versus *fer-4* mutant roots as indicated by qRT-PCR. **D-E**, *ERF114* and *ERF115* expression in root tips (**D**) or early stage LRP (**E**) under control conditions. **F-H,** LRP density of fer-4 mutants (**F**), and GUS staining (**G**) or confocal imaging (**H**) of *pERF115:NLS-GFP/GUS* root tips after 24h of BLM treatment. Scale bars= 50 μm. **I,** Quantification of GFP integrated density of BLM treated *pERF115:NLS-GFP/GUS* root tips

While these data suggested the possibility of a perceived mechanical or cell wall stress during the LR initiation, it cannot rule out the involvement of non-stress related processes activating *BZR1*. To address this shortcoming, we utilized a reverse genetics approach. It was previously suggested that FERONIA (FER), a member of RECEPTOR LIKE KINASE (RLK) family, works antagonistically to the BR signaling and negatively regulates cell extensibility (Hofte, 2015). FER has been shown to sense growth-related mechanical stress and activate compensatory, growth-limiting pathways. The loss of function mutant *fer-4* was characterized by an increased strain in the root elongation zone and an accelerated development of LRs (Shih *et al*., 2014). Therefore, we hypothesized that if a growth-related mechanical strain controls the activation of *ERF114* and *ERF115* expression, *fer-4* mutant seedlings might display increased levels of *ERF114* and *ERF115*. Indeed, the expression levels of both *ERF114* and *ERF115* were significantly increased in the 1-cm distal root tip section of *fer-4* mutants, compared to the wild type (Fig. 7C). To confirm this observation, we introduced the *ERF114* and *ERF115* transcriptional reporter lines into the *fer-4* background. Hyperactivation of *ERF114* and *ERF115* was clearly visible in aerial tissues, as indicated by intense GUS staining at the cotyledon-hypocotyl intersection (Supplementary Fig. 4A). In the roots, *ERF115* expression was strongly enhanced mainly in the protoxylem cell file (Fig. 7D). *ERF114* expression was also substantially increased in the pericycle cells and LR primordia (Fig. 7D,E). This coincided with a significantly increased LRP density in *fer-4* seedlings compared to the wild type (Fig. 7F). Treatment with auxin resulted in a visibly stronger induction of both *ERF114* and *ERF115* in the roots of the *fer-4* background compared to the wild type (Supplementary Fig. 4B,C), suggesting that FER might negatively regulate the auxin responsiveness of *ERF114* and *ERF115* expression.

Next, we investigated if FER might also restrain wound-induced *ERF115* expression, which was previously demonstrated to be accompanied by auxin accumulation (Canher *et al*., 2020). Induction of stem cell death after a 24-h treatment of of *pERF115:GFP-NLS/GUS* seedlings with BLM resulted in an enhanced induction of *ERF115* in the *fer-4* background (Fig. 7G). Quantification of GFP integrity density from confocal images revealed a 74% increase (p<0,001) in *fer-4* compared to the wild type (Fig. 7I). Taken together, these results suggest that developmental and wound-specific *ERF114* and *ERF115* expression might be influenced by growth-related mechanical cues, with FER-dependent cell wall signaling negatively regulating auxin sensitivity in a tissue-specific manner.

## DISCUSSION

### Novel developmental roles for *ERF114* and *ERF115* in xylem and LR formation

*ERF114* and *ERF115* have been the focus of extensive research regarding wound response and regeneration. However, presumably due to their low expression levels, involvement of ERF114 and ERF115 in a developmental context has not been explored besides regulation of QC cell divisions (Heyman *et al*., 2013). In this study, we demonstrated that ERF115 and ERF114 act redundantly during meristem regeneration following wounding. High levels of *ERF114* and *ERF115* are also associated with xylem formation, which is essential for establishing xylem reconnection during hypocotyl grafting. Moreover, we showed that *ERF114* and *ERF115* are inducers of LR development. Even though *ERF115* expression could only be detected in protoxylem, *erf115* mutants displayed a reduced LR formation and auxin maxima density. Smetana *et al*. showed that during secondary growth, immature xylem cells carrying local auxin maxima act non-cell autonomously as organizer cells. Clonal activation of auxin signaling via MP was sufficient to trigger xylem vessel differentiation and subsequent formative divisions in the adjacent procambial and pericycle cells (Smetana et al., 2019). Previously, misexpression of *ERF115* in the endodermis, in combination with auxin accumulation, was shown to non-cell autonomously produce formative divisions in columella stem cells (Canher *et al*., 2020). Therefore, based on its protoxylem-specific expression and being an upstream inducer of *MP*, ERF115 might act non-cell autonomously on the adjacent pericycle cells carrying an auxin maximum to grant stem cell identity to founder cells. *ERF114* expression can be detected in LR founder cells and its overexpression, unlike *ERF115*, led to the formation of LRs in the presence of NPA. Mehrnia *et al*. previously described ERF114 as an inducer of axillary bud formation and outgrowth, where *Pectin Methyl Esterase Super Family Protein* (At3g62820), auxin influx carrier *AUXIN RESISTANT 1* and *CYCLIN D3;3* were among the induced genes in an *ERF114*-inducible overexpression system (Mehrnia *et al*., 2013). Cell wall softening using external pectin methyl esterase application was sufficient to result in lateral organ initiation in auxin transport-deficient *pin1* mutants (Braybrook and Peaucelle, 2013). However, artificial induction of either cell cycle progression or cell wall softening were found to be insufficient for the formation of fully grown lateral organs (Braybrook and Peaucelle, 2013; Vanneste et al., 2005). Therefore, it can be speculated that ERF114 might enable LR growth despite the inhibition of polar auxin transport by a combination of activating the cell cycle, softening the cell wall, and promoting auxin influx. Taken together, it is likely that ERF114 and ERF115 might regulate different parts of the LR formation process by acting on different downstream targets.

### *ERF114* and *ERF115* expression during growth and regeneration is driven by mechanical cues and regulated antagonistically by FER and BR signaling

Time-lapse confocal imaging revealed a series of events displaying a spatial-temporal correlation, starting with *ERF114* induction in the LR founder cell, followed by DR5 activation in the founder cell and ending with the programmed cell death of the adjacent protoxylem cell expressing *ERF115.* We show that mechanical bending is sufficient to induce *ERF114*. Moreover, *fer-4* mutants, which are associated with an increased maximal strain and LR density, displayed a strong activation of both *ERF114* and *ERF115* (Dong et al., 2019). Application of auxin resulted in an additional hyperactivation of *ERF114* and *ERF115* in *fer-4* seedlings. Our results support a framework in which mechanical cues drive the expression of *ERF114* and *ERF115* during both regeneration and development. In this framework, cell wall integrity and mechanosensory FERONIA-mediated signaling constitutes a feedback mechanism that negatively regulate the expression of *ERF114* and *ERF115* in LR and xylem cells, respectively. In the absence of wounding, developmentally driven mechanical pressures drive low levels of *ERF114* and *ERF115* expression. Reciprocal activation of MP-mediated auxin signaling by *ERF115* in turn likely drives xylem formation and LR development. While activation of *ERF114* by mechanical strain promotes LR progression and outgrowth, the exact downstream targets potentially unique to ERF114 are yet to be identified. In the context of regeneration, wounding and the sudden cell collapse following cell death would create a perturbation in mechanical homeostasis larger than that resulting from normal developmental activity. The inability of the gradual FER-mediated antagonistic feedback to re-instate the mechanical homeostasis would lead to the strong and robust *ERF114* and *ERF115* expression observed, which in turn activates developmental pathways for successful regeneration.

The mechanisms that cause the induction of *ERF114* and *ERF115* upon mechanical cues still need to be investigated. Several CWI sensors associated with mechano-perception such as members of the MECHANOSENSITIVE CHANNEL OF SMALL CONDUCTANCE LIKE (MSL) and MID1-COMPLEMENTING ACTIVITY A (MCA) gene families are prime candidates (Bacete and Hamann, 2020). Additionally, the cell wall integrity sensor RECEPTOR LIKE PROTEIN 44 (RLP44) stands out as a putative upstream regulator, as it was shown to mediate the activation of BR signaling in response to cell wall damage (Wolf et al., 2014). Furthermore, RLP44 was demonstrated to regulate phytosulfokine signaling through its interaction partner PHYTOSULFOKINE RECEPTOR 1 (PSKR1) and prevent xylem fate acquirement in the cambium (Holzwart et al., 2018). Considering that ERF115 is a direct activator of *PSK5*, RLP44 is worth investigation as a wall integrity sensor activating *ERF115* and *PSK5* expression during development and regeneration.

## METHODS

### Plant materials and growth conditions

Plants were grown under a long-day regime (16-h light/8-h darkness) on agar-solidified culture medium (Murashige and Skoog [MS] medium, 10 g/l saccharose, 4.3 g/l 2-(N-morpholino) ethanesulfonic acid [MES], and 0.8% plant tissue culture agar) at 21°C. MP:MP-GFP seeds were kindly provided by Dolf Weijers, Wageningen University, Netherlands (Cole et al., 2009; Schlereth et al., 2010). *pPASPA3:tdTomato-NLS* (Xuan et al., 2016), *DR5:LUC* (Moreno-Risueno *et al*., 2010), *ERF115^OE^*, *ERF115^SRDX^*, *pERF115:NLS-GFP/GUS* and *erf115* (SALK_021981) were described previously (Heyman *et al*., 2013). The *pERF114-NLS-GFP/GUS* reporter was created by cloning the 2163 nucleotide promoter region upstream of *ERF114* start codon into the pCR™Blunt II- TOPO® vector. The resulting entry vector was cloned into pMK7S*NFm14GW (Karimi et al., 2002) and transformed into the Col-0 background by agrobacterium-mediated transformation. The *ERF114* CRISPR mutant (Supplementary Fig. 6) was created using a dual guide RNA approach (Pauwels et al., 2018). A construct targeting two different sites in the *ERF114* gene was designed where the single guide RNA1 (sgRNA1, pMR217_pDONR_P1P) and sgRNA2 (pMR218_pDONR_P5P) were annealed and inserted via a cut ligation method using Bbs1 in pMR217_pDONR_P1P and pMR218_pDONR_P5P, respectively. Next, using a Gateway LR reaction the two sgRNAs were combined into pDE_CAS9_Basta to yield the final expression clone. The *erf114 115* double mutants (Supplementary Fig. 5) were obtained by transforming the *ERF11*4 CRSIPR construct into the *erf115* single mutant by floral dip method. Primary transformants were selected on agar plates containing Basta and genotyped for the deleted fragment and further confirmed using Sanger sequencing. Cas9 free and homozygous mutants were selected in the T3 generation for further use. Primers used for the CRISPR-mediated mutagenesis and genotyping are described in Supplementary Table 1.

### Lateral root staging

Eight-day-old seedlings were fixed in 80% acetone overnight. After discarding the acetone, seedlings were treated with Clearsee optical clearing solution overnight to increase transparency of tissues (Kurihara et al., 2015). The next day, Clearsee solution was discarded, and seedlings were rinsed with distilled water to wash off the remaining Clearsee solution. Further clearing and staging was performed as described previously, starting with the acid clearing step (Malamy, 1997). The counting of the LRs was done using an Olympus BX51 DIC microscope.

### DR5:LUC imaging

A Lumazone imaging system equipped with a charge-coupled device (CCD) camera (Princeton Instruments, Trenton, NJ, USA) was used for luciferase imaging. The CCD camera is controlled by WinView/32 software. For imaging DR5:LUC expression, square plates containing ½ MS medium with or without chemicals were sprayed with 1-mM D-Luciferin solution (0.01% Tween80) and left to dry in the dark. Eight-day-old DR5:LUC seedlings were transferred onto the plates and imaged immediately with a macro lens with a 20-min exposure time for indicated time points. For time-lapse imaging, an image was acquired every 20 min. The picture series were saved as TIFF format by WinView/32 software for further analysis in ImageJ (http://imagej.nih.gov/ij/). The prebranch site densities were calculated by dividing the number of DR5 maxima by the root length. The gravitational bending assay was done by rotating the square plate holding the seedlings 90° and imaging the *DR5:LUC* overnight with 30-min intervals. Number of *DR5:LUC* maxima at the bend site was calculated from the obtained time-lapse imaging.

### Histochemical assays

β-glucuronidase (GUS) staining was performed as described previously (Lammens et al., 2008).

### Quantitative RT-PCR

RNA was isolated from the respective tissues with the RNeasy isolation kit (Qiagen). DNase treatment with the RQ1 RNase-Free DNase (Promega) was performed before cDNA synthesis with the iScript cDNA Synthesis Kit (Bio-Rad). Relative expression levels were determined by qRT-PCR with the LightCycler 480 Real-Time SYBR Green PCR System (Roche). The primers used are described in Supplementary Table 2. The *RPS26C* and *EMB238*6 reference genes were used for normalization. In three biological repetitions, total RNA was isolated by means of the RNeasy Plant mini kit (Qiagen). For the root tips, seedlings were sown and grown for 5 days on nylon meshes (Prosep) and subsequently harvested using a scalpel. Quantitative PCR data were analyzed using the 2(–ΔΔCt) method.

### Statistical analysis

Statistical analysis was performed using SAS Enterprise Guide 7.15 HF8. For LR quantifications, the mixed model procedure was used to calculate significance values for pairwise comparisons with the wild type by least squares post-hoc tests and corrected for multiple testing using the Bonferroni Method. Genotype was added as a main effect and experimental repeats were included as random effect. Degrees of freedom were calculated by the Kenward-Rogers method. For regeneration assays probabilities of successful regeneration were compared using logistic regression analysis. Odds ratios with 99% Wald confidence limits and p-values were calculated after Fisher’s exact testing.

### Regeneration assays

Grafting was performed as described previously and xylem connectivity was determined by CFDA fluorescence in the cotyledons (Melnyk, 2017). For assessment of xylem bridge formation in LRs, 3-day-old seedlings were rotated 90° to synchronously induce LRs. Stringent root tip cutting was performed at a 250-µM distance from the root tip, as described previously (Sena et al., 2009).

### Treatments

For germination for the NPA experiments, 125mM NPA stock in DMSO was diluted to a final concentration of 10 μM in ½ MS agar medium. Seedlings were germinated and grown on NPA medium for indicated times, stained with Lugol solution and imaged by light microscopy. Basic Fuchsin staining in combination with Clearsee was performed as described previously (Ursache et al., 2018). For BLM treatments, 5-day-old seedlings were transferred to medium supplemented with 0.6 mg/L bleomycin sulphate (Calbiochem) for 24 h.

### Confocal and light microscopy

Arabidopsis roots were stained using PI by incubation in a 10-μM solution for 3 min before imaging. Imaging and laser ablation was performed on a Leica TCS SP8 X microscope equipped with an Argon laser (488 nm) for GFP excitation and a white light laser (554 nm) for tdTomato excitation. GFP and tdTomato emissions were collected at 500-540 nm and 570-630 nm, respectively. Time-lapse imaging was performed by acquiring multiple z-slice images of seedlings mounted in live imaging chambers every 30 min. Manual root bending was performed as described previously (Ditengou *et al*., 2008). Leica LAS X and Fiji were used for further image processing. For light and differential interference contrast microscopy, an Olympus BX51 microscope was used.

## SUPPLEMENTAL INFORMATION

Supplemental Information is available at Molecular Plant Online

## AUTHOR CONTRIBUTIONS

B.C. and L.D.V. wrote the manuscript. F.L, F.A and B.C performed LR phenotyping. BC performed luminescence imaging and phenotyping. A.Z, S.M and C.W.M. performed the grafting assays. J.H and A.B performed CRISPR-Cas9 mutagenesis. S.W. analyzed vascular development. All authors approved the manuscript.

## ACKNOWLEDGEMENTS

The authors thank Annick Bleys for help in preparing the manuscript. This work was supported by grants (G007218N and G010820N) and a predoctoral fellowship (F.L.) from the Research Foundation-Flanders. AZ, SM and CWM were supported by a Wallenberg Academy Fellowship (2016-0274) and a Vetenskapsrådet grant (2017-05122).

## REFERENCES

1. Bhatia, N., Bozorg, B., Larsson, A., Ohno, C., Jonsson, H., and Heisler, M.G. (2016). Auxin Acts through MONOPTEROS to Regulate Plant Cell Polarity and Pattern Phyllotaxis. Curr Biol 26:3202–3208. 10.1016/j.cub.2016.09.044.

2. Birnbaum, K.D., and Sanchez Alvarado, A. (2008). Slicing across kingdoms: regeneration in plants and animals. Cell 132:697–710. 10.1016/j.cell.2008.01.040.

3. Braybrook, S.A., and Peaucelle, A. (2013). Mechano-chemical aspects of organ formation in Arabidopsis thaliana: the relationship between auxin and pectin. PLoS One 8:e57813. 10.1371/journal.pone.0057813.

4. Canher, B., Heyman, J., Savina, M., Devendran, A., Eekhout, T., Vercauteren, I., Prinsen, E., Matosevich, R., Xu, J., Mironova, V., et al. (2020). Rocks in the auxin stream: Wound-induced auxin accumulation and ERF115 expression synergistically drive stem cell regeneration. Proc Natl Acad Sci U S A 117:16667–16677. 10.1073/pnas.2006620117.

5. Casimiro, I., Marchant, A., Bhalerao, R.P., Beeckman, T., Dhooge, S., Swarup, R., Graham, N., Inze, D., Sandberg, G., Casero, P.J., et al. (2001). Auxin transport promotes Arabidopsis lateral root initiation. Plant Cell 13:843–852. 10.1105/tpc.13.4.843.

6. Cole, M., Chandler, J., Weijers, D., Jacobs, B., Comelli, P., and Werr, W. (2009). DORNROSCHEN is a direct target of the auxin response factor MONOPTEROS in the Arabidopsis embryo. Development 136:1643–1651. 10.1242/dev.032177.

7. De Smet, I. (2010). Multimodular auxin response controls lateral root development in Arabidopsis. Plant Signal Behav 5:580–582. 10.4161/psb.11495.

8. De Smet, I., Tetsumura, T., De Rybel, B., Frei dit Frey, N., Laplaze, L., Casimiro, I., Swarup, R., Naudts, M., Vanneste, S., Audenaert, D., et al. (2007). Auxin-dependent regulation of lateral root positioning in the basal meristem of Arabidopsis. Development 134:681–690. 10.1242/dev.02753.

9. De Smet, I., Lau, S., Voss, U., Vanneste, S., Benjamins, R., Rademacher, E.H., Schlereth, A., De Rybel, B., Vassileva, V., Grunewald, W., et al. (2010). Bimodular auxin response controls organogenesis in Arabidopsis. Proc Natl Acad Sci U S A 107:2705–2710. 10.1073/pnas.0915001107.

10. Ditengou, F.A., Teale, W.D., Kochersperger, P., Flittner, K.A., Kneuper, I., van der Graaff, E., Nziengui, H., Pinosa, F., Li, X., Nitschke, R., et al. (2008). Mechanical induction of lateral root initiation in Arabidopsis thaliana. Proc Natl Acad Sci U S A 105:18818–18823. 10.1073/pnas.0807814105.

11. Dong, Q., Zhang, Z., Liu, Y., Tao, L.Z., and Liu, H. (2019). FERONIA regulates auxin-mediated lateral root development and primary root gravitropism. FEBS Lett 593:97–106. 10.1002/1873-3468.13292.

12. Fendrych, M., Van Hautegem, T., Van Durme, M., Olvera-Carrillo, Y., Huysmans, M., Karimi, M., Lippens, S., Guerin, C.J., Krebs, M., Schumacher, K., et al. (2014). Programmed cell death controlled by ANAC033/SOMBRERO determines root cap organ size in Arabidopsis. Curr Biol 24:931–940. 10.1016/j.cub.2014.03.025.

13. Feng, W., Kita, D., Peaucelle, A., Cartwright, H.N., Doan, V., Duan, Q., Liu, M.C., Maman, J., Steinhorst, L., Schmitz-Thom, I., et al. (2018). The FERONIA Receptor Kinase Maintains Cell-Wall Integrity during Salt Stress through Ca(2+) Signaling. Curr Biol 28:666–675 e665. 10.1016/j.cub.2018.01.023.

14. Franck, C.M., Westermann, J., and Boisson-Dernier, A. (2018). Plant Malectin-Like Receptor Kinases: From Cell Wall Integrity to Immunity and Beyond. Annu Rev Plant Biol 69:301–328. 10.1146/annurev-arplant-042817-040557.

15. Heyman, J., Canher, B., Bisht, A., Christiaens, F., and De Veylder, L. (2018). Emerging role of the plant ERF transcription factors in coordinating wound defense responses and repair. J Cell Sci 131 10.1242/jcs.208215.

16. Heyman, J., Cools, T., Vandenbussche, F., Heyndrickx, K.S., Van Leene, J., Vercauteren, I., Vanderauwera, S., Vandepoele, K., De Jaeger, G., Van Der Straeten, D., et al. (2013). ERF115 controls root quiescent center cell division and stem cell replenishment. Science 342:860–863. 10.1126/science.1240667.

17. Heyman, J., Cools, T., Canher, B., Shavialenka, S., Traas, J., Vercauteren, I., Van den Daele, H., Persiau, G., De Jaeger, G., Sugimoto, K., et al. (2016). The heterodimeric transcription factor complex ERF115-PAT1 grants regeneration competence. Nat Plants 2:16165. 10.1038/nplants.2016.165.

18. Article I. Hoermayer, L., Montesinos, J.C., Marhava, P., Benkova, E., Yoshida, S., and Friml, J. (2020). Wounding-induced changes in cellular pressure and localized auxin signalling spatially coordinate restorative divisions in roots. Proc Natl Acad Sci U S A 117:15322-15331. 10.1073/pnas.2003346117.

19. Hofte, H. (2015). The yin and yang of cell wall integrity control: brassinosteroid and FERONIA signaling. Plant Cell Physiol 56:224–231. 10.1093/pcp/pcu182.

20. Hofte, H., and Voxeur, A. (2017). Plant cell walls. Curr Biol 27:R865–R870. 10.1016/j.cub.2017.05.025.

21. Holzwart, E., Huerta, A.I., Glockner, N., Garnelo Gomez, B., Wanke, F., Augustin, S., Askani, J.C., Schurholz, A.K., Harter, K., and Wolf, S. (2018). BRI1 controls vascular cell fate in the Arabidopsis root through RLP44 and phytosulfokine signaling. Proc Natl Acad Sci U S A 115:11838–11843. 10.1073/pnas.1814434115.

22. Jürgens, T.B.a.G. (1993). The role of the monopteros gene in organising the basal body region of the Arabidopsis embryo. Development 118:575–587.

23. Karimi, M., Inze, D., and Depicker, A. (2002). GATEWAY vectors for Agrobacterium-mediated plant transformation. Trends Plant Sci 7:193–195. 10.1016/s1360-1385(02)02251-3.

24. Kong, X., Tian, H., Yu, Q., Zhang, F., Wang, R., Gao, S., Xu, W., Liu, J., Shani, E., Fu, C., et al. (2018). PHB3 Maintains Root Stem Cell Niche Identity through ROS-Responsive AP2/ERF Transcription Factors in Arabidopsis. Cell Rep 22:1350–1363. 10.1016/j.celrep.2017.12.105.

25. Kurihara, D., Mizuta, Y., Sato, Y., and Higashiyama, T. (2015). ClearSee: a rapid optical clearing reagent for whole-plant fluorescence imaging. Development 142:4168–4179. 10.1242/dev.127613.

26. Lammens, T., Boudolf, V., Kheibarshekan, L., Zalmas, L.P., Gaamouche, T., Maes, S., Vanstraelen, M., Kondorosi, E., La Thangue, N.B., Govaerts, W., et al. (2008). Atypical E2F activity restrains APC/CCCS52A2 function obligatory for endocycle onset. Proc Natl Acad Sci U S A 105:14721–14726. 10.1073/pnas.0806510105.

27. Article II. Lee, H.S., Kim, Y., Pham, G., Kim, J.W., Song, J.H., Lee, Y., Hwang, Y.S., Roux, S.J., and Kim, S.H. (2015). Brassinazole resistant 1 (BZR1)-dependent brassinosteroid signalling pathway leads to ectopic activation of quiescent cell division and suppresses columella stem cell differentiation. J Exp Bot 66:4835- 4849. 10.1093/jxb/erv316.

28. Lucas, M., Godin, C., Jay-Allemand, C., and Laplaze, L. (2008). Auxin fluxes in the root apex co-regulate gravitropism and lateral root initiation. J Exp Bot 59:55–66. 10.1093/jxb/erm171.

29. Luo, L., Zeng, J., Wu, H., Tian, Z., and Zhao, Z. (2018). A Molecular Framework for Auxin-Controlled Homeostasis of Shoot Stem Cells in Arabidopsis. Mol Plant 11:899–913. 10.1016/j.molp.2018.04.006.

30. Malamy, J.E., and Benfey, P. N. (1997). Organization and cell differentiation in lateral roots of Arabidopsis thaliana. Development 124:33–44.

31. Marhava, P., Hoermayer, L., Yoshida, S., Marhavy, P., Benkova, E., and Friml, J. (2019). Re-activation of Stem Cell Pathways for Pattern Restoration in Plant Wound Healing. Cell 177:957–969 e913. 10.1016/j.cell.2019.04.015.

32. Marhavy, P., Montesinos, J.C., Abuzeineh, A., Van Damme, D., Vermeer, J.E., Duclercq, J., Rakusova, H., Novakova, P., Friml, J., Geldner, N., et al. (2016). Targeted cell elimination reveals an auxin-guided biphasic mode of lateral root initiation. Genes Dev 30:471–483. 10.1101/gad.276964.115.

33. Mehrnia, M., Balazadeh, S., Zanor, M.I., and Mueller-Roeber, B. (2013). EBE, an AP2/ERF transcription factor highly expressed in proliferating cells, affects shoot architecture in Arabidopsis. Plant Physiol 162:842–857. 10.1104/pp.113.214049.

34. Melnyk, C.W. (2017). Connecting the plant vasculature to friend or foe. New Phytol 213:1611–1617. 10.1111/nph.14218.

35. Melnyk, C.W., Schuster, C., Leyser, O., and Meyerowitz, E.M. (2015). A Developmental Framework for Graft Formation and Vascular Reconnection in Arabidopsis thaliana. Curr Biol 25:1306–1318. 10.1016/j.cub.2015.03.032.

36. Melnyk, C.W., Gabel, A., Hardcastle, T.J., Robinson, S., Miyashima, S., Grosse, I., and Meyerowitz, E.M. (2018). Transcriptome dynamics at Arabidopsis graft junctions reveal an intertissue recognition mechanism that activates vascular regeneration. Proc Natl Acad Sci U S A 115:E2447–E2456. 10.1073/pnas.1718263115.

37. Moller, B.K., Xuan, W., and Beeckman, T. (2017). Dynamic control of lateral root positioning. Curr Opin Plant Biol 35:1–7. 10.1016/j.pbi.2016.09.001.

38. Moreno-Risueno, M.A., Van Norman, J.M., Moreno, A., Zhang, J., Ahnert, S.E., and Benfey, P.N. (2010). Oscillating Gene Expression Determines Competence for Periodic *Arabidopsis* Root Branching. Science 329:1306–1311. 10.1126/science.1191937.

39. Parizot, B., Laplaze, L., Ricaud, L., Boucheron-Dubuisson, E., Bayle, V., Bonke, M., De Smet, I., Poethig, S.R., Helariutta, Y., Haseloff, J., et al. (2008). Diarch symmetry of the vascular bundle in Arabidopsis root encompasses the pericycle and is reflected in distich lateral root initiation. Plant Physiol 146:140–148. 10.1104/pp.107.107870.

40. Pauwels, L., De Clercq, R., Goossens, J., Inigo, S., Williams, C., Ron, M., Britt, A., and Goossens, A. (2018). A Dual sgRNA Approach for Functional Genomics in Arabidopsis thaliana. G3 (Bethesda) 8:2603–2615. 10.1534/g3.118.200046.

41. Perianez-Rodriguez, J., Manzano, C., and Moreno-Risueno, M.A. (2014). Post-embryonic organogenesis and plant regeneration from tissues: two sides of the same coin? Front Plant Sci 5:219. 10.3389/fpls.2014.00219.

42. Przemeck, G.K., Mattsson, J., Hardtke, C.S., Sung, Z.R., and Berleth, T. (1996). Studies on the role of the Arabidopsis gene MONOPTEROS in vascular development and plant cell axialization. Planta 200:229–237. 10.1007/bf00208313.

43. Savatin, D.V., Gramegna, G., Modesti, V., and Cervone, F. (2014). Wounding in the plant tissue: the defense of a dangerous passage. Front Plant Sci 5:470. 10.3389/fpls.2014.00470.

44. Schlereth, A., Moller, B., Liu, W., Kientz, M., Flipse, J., Rademacher, E.H., Schmid, M., Jurgens, G., and Weijers, D. (2010). MONOPTEROS controls embryonic root initiation by regulating a mobile transcription factor. Nature 464:913–916. 10.1038/nature08836.

45. Sena, G., Wang, X., Liu, H.Y., Hofhuis, H., and Birnbaum, K.D. (2009). Organ regeneration does not require a functional stem cell niche in plants. Nature 457:1150–1153. 10.1038/nature07597.

46. Shih, H.W., Miller, N.D., Dai, C., Spalding, E.P., and Monshausen, G.B. (2014). The receptor-like kinase FERONIA is required for mechanical signal transduction in Arabidopsis seedlings. Curr Biol 24:1887–1892. 10.1016/j.cub.2014.06.064.

47. Smetana, O., Makila, R., Lyu, M., Amiryousefi, A., Sanchez Rodriguez, F., Wu, M.F., Sole-Gil, A., Leal Gavarron, M., Siligato, R., Miyashima, S., et al. (2019). High levels of auxin signalling define the stem-cell organizer of the vascular cambium. Nature 565:485–489. 10.1038/s41586-018-0837-0.

48. Sun, S.B., Meng, L.S., Sun, X.D., and Feng, Z.H. (2010). Using high competent shoot apical meristems of cockscomb as explants for studying function of ASYMMETRIC LEAVES2-LIKE11 (ASL11) gene of Arabidopsis. Mol Biol Rep 37:3973–3982. 10.1007/s11033-010-0056-8.

49. Ursache, R., Andersen, T.G., Marhavy, P., and Geldner, N. (2018). A protocol for combining fluorescent proteins with histological stains for diverse cell wall components. Plant J 93:399–412. 10.1111/tpj.13784.

50. Vanneste, S., De Rybel, B., Beemster, G.T., Ljung, K., De Smet, I., Van Isterdael, G., Naudts, M., Iida, R., Gruissem, W., Tasaka, M., et al. (2005). Cell cycle progression in the pericycle is not sufficient for SOLITARY ROOT/IAA14- mediated lateral root initiation in Arabidopsis thaliana. Plant Cell 17:3035–3050. 10.1105/tpc.105.035493.

51. Vukasinovic, N., Wang, Y., Vanhoutte, I., Fendrych, M., Guo, B., Kvasnica, M., Jiroutova, P., Oklestkova, J., Strnad, M., and Russinova, E. (2021). Local brassinosteroid biosynthesis enables optimal root growth. Nat Plants 7:619–632. 10.1038/s41477-021-00917-x.

52. Wolf, S., Mravec, J., Greiner, S., Mouille, G., and Höfte, H. (2012a). Plant cell wall homeostasis is mediated by brassinosteroid feedback signaling. Current Biology 22:1732–1737. 10.1016/j.cub.2012.07.036.

53. Wolf, S., Mravec, J., Greiner, S., Mouille, G., and Hofte, H. (2012b). Plant cell wall homeostasis is mediated by brassinosteroid feedback signaling. Curr Biol 22:1732–1737. 10.1016/j.cub.2012.07.036.

54. Wolf, S., van der Does, D., Ladwig, F., Sticht, C., Kolbeck, A., Schurholz, A.K., Augustin, S., Keinath, N., Rausch, T., Greiner, S., et al. (2014). A receptor-like protein mediates the response to pectin modification by activating brassinosteroid signaling. Proc Natl Acad Sci U S A 111:15261–15266. 10.1073/pnas.1322979111.

55. Xuan, W., Band, L.R., Kumpf, R.P., Van Damme, D., Parizot, B., De Rop, G., Opdenacker, D., Möller, B.K., Skorzinski, N., Njo, M.F., et al. (2016). Cyclic programmed cell death stimulates hormone signaling and root development in Arabidopsis Science 351:384–387. 10.1126/science.aad2776.

56. Zhou, W., Lozano-Torres, J.L., Blilou, I., Zhang, X., Zhai, Q., Smant, G., Li, C., and Scheres, B. (2019). A Jasmonate Signaling Network Activates Root Stem Cells and Promotes Regeneration. Cell 177:942–956 e914. 10.1016/j.cell.2019.03.006.

