## Supplemental Figures and Tables for "The regeneration factors ERF114 and ERF115 act as transducers of mechanical cues to developmental pathways"

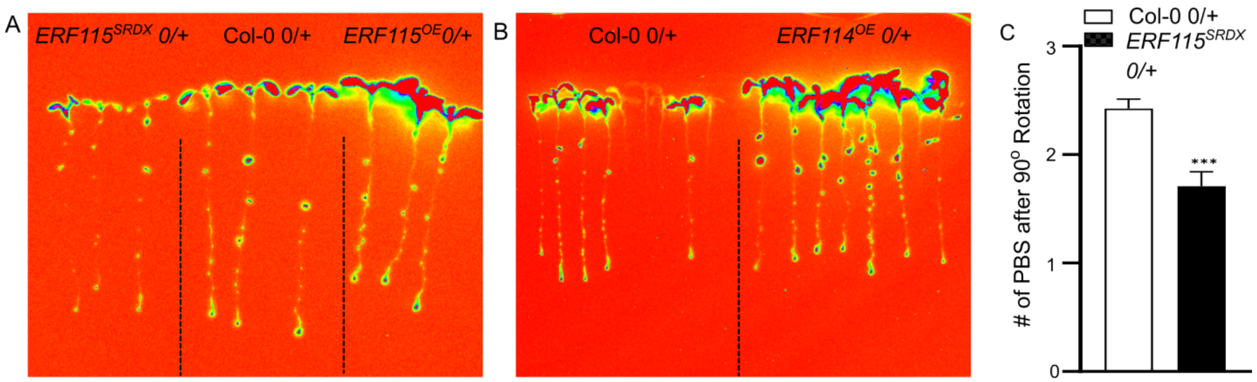


**Supplementary Fig. 1. ERF115 enhances the auxin maxima at lateral root initiation sites** (A,B) Representative luminescence images of hemizygous F1 seedlings of *DR5:LUC* crossed with indicated genotypes. Brightness and contrast settings were adjusted for better visualization. (C) Number of *DR5:LUC* maxima formed at the bend site following 90° rotation (***p<0.001, Students T-test).

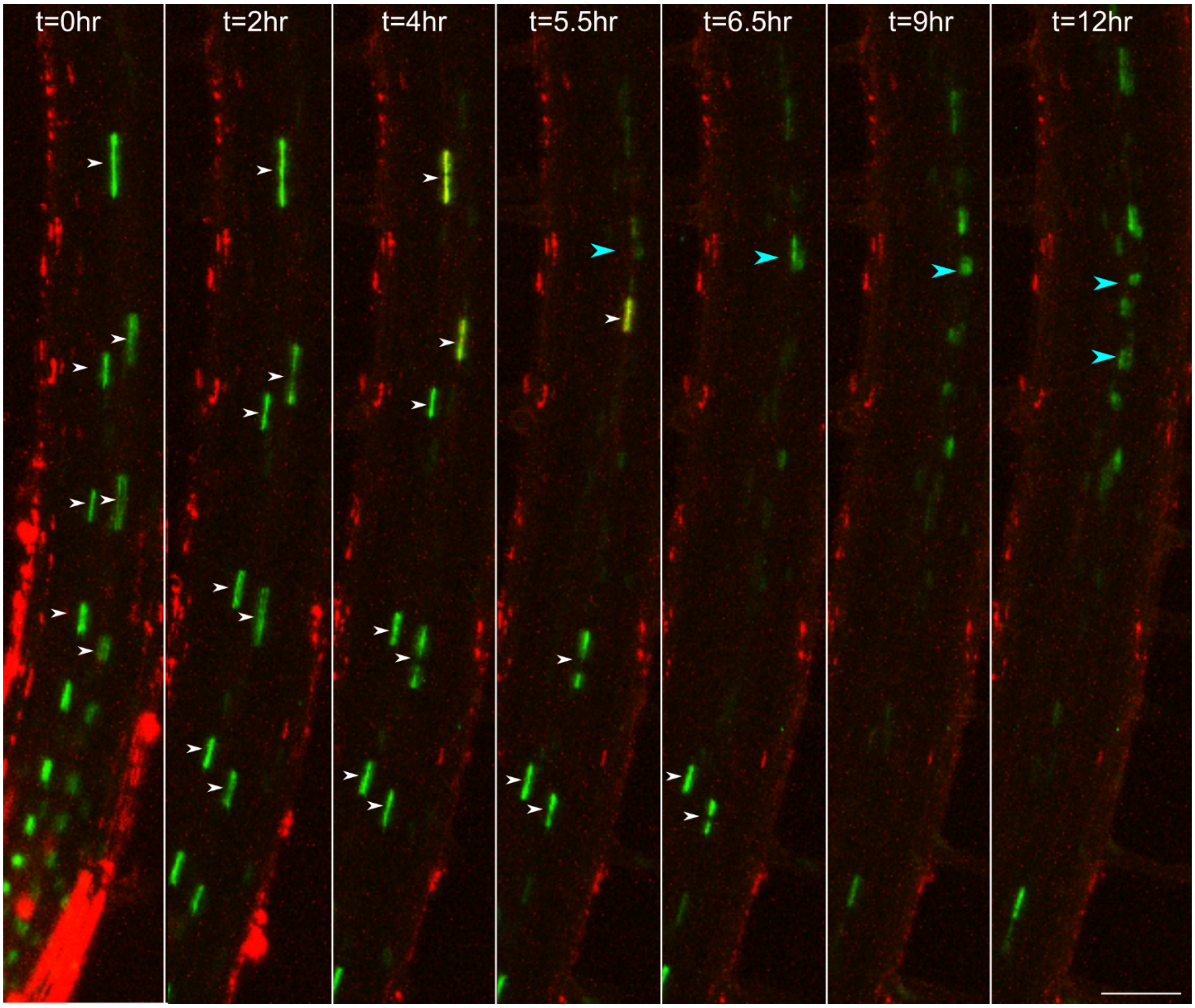


**Supplementary Fig. 2. Stills from time-lapse imaging of *pDR5:Venus-NLS pPASPA3:tdTomato-NLS* dual reporter FC formation (Supplementary Movie 3).** White arrowheads point to *DR5:VENUS-NLS* positive protoxylem nuclei. Yellow colored nuclei contain both *DR5:Venus-NLS* and *PASPA3:tdTomato-NLS* signals. Blue arrowheads point to LRP nuclei marked by *DR5:VENUS-NLS* fluorescence. Scalebar = 50 μm.

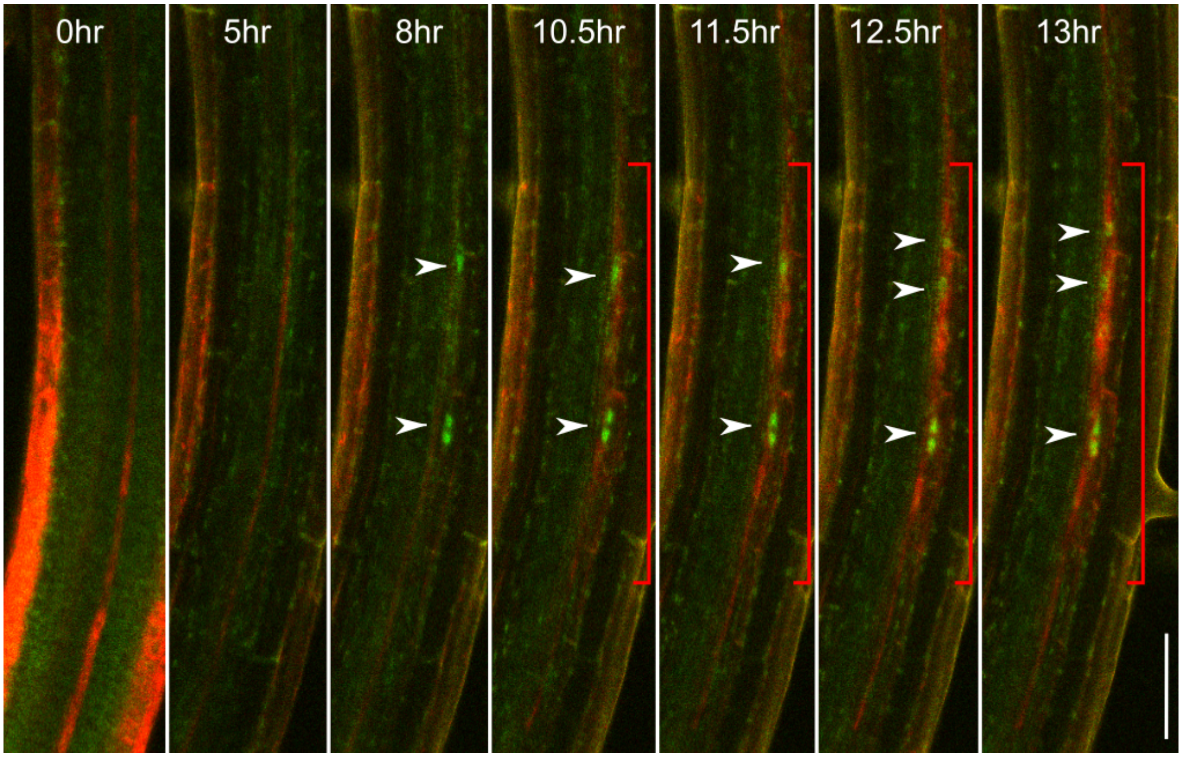


**Supplementary Fig. 3. Stills from time-lapse confocal imaging of *pERF114:GFP-NLS/GUS pDR5:RFP* during LR formation (Supplementary Movie 4).** Arrowheads point to a nuclear GFP signal. Red bar indicates the region of *DR5:RFP* signal accumulation Scale bar = 50μM.

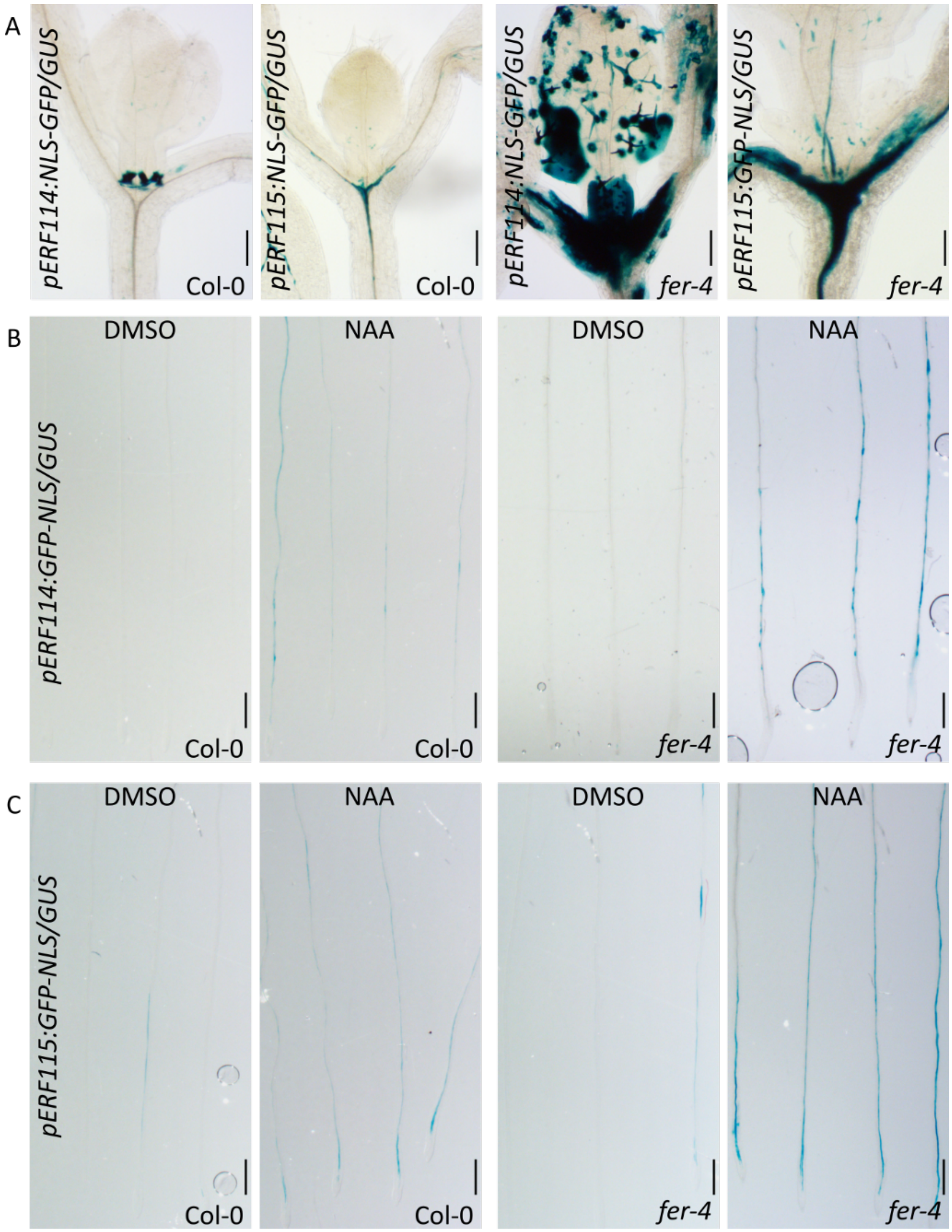


**Supplementary Fig. 4. *fer*-4 mutants display increased ERF114 and ERF115 expression that can be enhanced in combination with auxin** treatment (A) Light microscopy images of GUS-stained *pERF114:GFP-NLS/GUS* and *pERF115:GFP-NLS/GUS* seedlings showing a stronger expression in the *fer-4* background compared to Col-0. (B,C) GUS staining of the roots of seedlings showing a stronger expression of *ERF114* (B) and *ERF115* (C) after treatment with NAA in the *fer-4* background compared to the wild type (Col-0). Scale bars = 100 µM.

***erf114* single mutant**

-24 bp ATG

265 bp

GGAAGAAAAGGCTATATTTAAAAGAA ——— ACCAAGGTATATATATACAT (Col-WT)

GGAAGAAAAGGCTATATCAAATTTGC ......... ACCAAGGTATATATATACAT (*erf114*)

***erf114,115* double mutant**

+18 bp ATG

235 bp

GCAAAAGTAGATCCTCGGATGTATGGGAAGAGGCCT ——— ATATACATGCTCATATCG (Col-WT)

GCAAAAGTAGATCCTCGGATGTATGGGAAGAGGCTC ......... ATATACATGCTCATATCG (*erf114erf115*)

**Supplementary Fig. 5.** Outline of deletion events in *erf114* single and *erf114 115* double mutants. Red letters denote mismatched nucleotides.

**Supplementary Table 1.** Primers used in CRISPR-mediated mutagenesis and genotyping

| |
|---|
| **Guide RNAs (gRNAs)** |
| gRNA_1 : ATGTATGGGAAGAGGCCTTT |
| gRNA_2 : GCGCCTACTCATCAAGACCA |
| **Primers for genotyping *erf114* CRISPR and *erf114 115 mutants*** |
| erf114_F : CAATCTAATTTCTTCTCCTT |
| erf114_R : CGGTTTCAAATGTCCCGAGCC |

**Supplementary Table 2.** Primers used for qRT-PCR analysis

| | |
|---|---|
| qERF114-F | CGGAAACCAACAAAGCAAAT |
| qERF114_R | GGCCTTTGCCTTACACCTCT |
| qERF115-F | GGAAACCAAAGCAGCTCTCA |
| qERF115-R | GCAGCTTCAGCAGTCTCAAA |
| qRPS26C-F | GACTTTCAAGCGCAGGAATGGTG |
| qRPS26C-R | CCTTGTCCTTGGGGCAACACTTT |
| qEMB2386-F | CTCTCGTTCCAGAGCTCGCAAAA |
| qEMB2386-R | AAGAACACGCATCCTACGCATCC |

**Supplementary Movie 1.** Time-lapse confocal imaging of the *pERF115:NLS-GFP/GUS pPASPA3:tdTom-NLS* dual reporter line. Arrowheads indicate the protoxylem nuclei holding GFP and tdTom signals that disappear simultaneously, indicating the final nuclear disintegration stage of protoxylem maturation.

**Supplementary Movie 2.** Time-lapse confocal imaging of the *pERF115:NLS-GFP/GUS pDR5:RFP* dual reporter line. White arrowheads indicate GFP positive protoxylem nuclei residing inside the protoxylem cell carrying a RFP signal. Blue arrowheads indicate the accumulation of an RFP signal in the LR pre-branch site.

**Supplementary Movie 3.** Time-lapse confocal imaging of the *DR5:VENUS-NLS pPASPA3:tdTom-NLS* dual reporter line. Appearance of the nuclear VENUS signal (*DR5*) in the pericycle cells is succeeded by gradual accumulation of the RFP signal (*PASPA3*).

**Supplementary Movie 4.** Time-lapse confocal imaging of the *pERF114:NLS-GFP/GUS pDR5:RFP* dual reporter line. White arrowheads indicate GFP positive protoxylem nuclei residing inside the protoxylem cell carrying a RFP signal. Blue arrowheads indicate the accumulation of an RFP signal in the LR pre-branch site.

**Supplementary Movie 5.** Time-lapse confocal imaging of a PI-stained (red) *pERF114:NLS-GFP/GUS* reporter seedling 2 h after laserinduced cell wall damaging at the protoxylem cell boundary. The red stained strands indicate the spiral secondary cell wall structure of protoxylem. ERF114 induction in the pericycle is followed by anticlinal divisions.

**Supplementary Movie 6.** Time-lapse confocal imaging of a PI-stained (red) *pERF114:NLS-GFP/GUS pDR5:RFP* dual reporter seedling 1 h following manual bending. GFP and PI were excited with a 488-nM laser while RFP was excited with a 559-nm laser. GFP emission (green) was collected between 500-535 nm. PI emission (cyan) was collected between 575-675 nm and RFP emission (red) was collected between 550-560 nm.

**Supplementary Movie 7.** Time-lapse confocal imaging of the *pERF115:NLS-GFP/GUS pDR5:RFP* dual reporter line including the transmitted light channel 1 hour following manual bending. ERF115 induction in cells below the initiating LR primordia can be observed through appearance of a nuclear GFP signal.
